# Evolution of plasmid domestication in plant-associated *Pantoea*: massive gain of genetic redundancy followed by differential gene loss

**DOI:** 10.1101/2024.10.17.618910

**Authors:** Devani Romero Picazo, Paul Kwasigroch, Nils F. Hülter, Tal Dagan

## Abstract

Plasmids are important drivers of evolutionary transformations and ecological adaptation in prokaryotes. Plasmids supplying the host with beneficial functions may become domesticated and gain a stable inheritance within the host lineage in the form of secondary chromosomes. Chromosomes descendent from plasmids (termed chromids) comprise core genes that are universally present in all taxon members. The origin of plasmid core genes remains poorly understood and alternative hypotheses entailing gene translocation or genetic redundancy are debated. Studying plasmid evolution in plant-associated *Pantoea*, we show that the large *Pantoea* plasmid (LPP) and plasmid pPag1 comprise core genes in the host taxa. We infer the LPP acquisition in plant-associated ancestors, while the pPag1 acquisition is traced to the ancestor of plant growth-promoting species. We show that both plasmids are vertically inherited and the LPP replication is furthermore coordinated with cell division, hence they are better described as chromids. Using phylogenomics we infer that the plasmid acquisition manifested in massive gene gain, some redundant with the chromosome, followed by differential gene loss. Our results provide an explanation for the evolution of core plasmid genes that entails plasmid-driven species divergence, which preliminarily manifests in genetic redundancy, and is followed by gradual loss of redundant genes.

**Significance statement:** Adaptation to host-associated lifestyle in bacteria can be promoted by the acquisition of plasmids, which are mobile extra-chromosomal genetic elements. A canonic example is the symbiosis *Rhizobia* plasmids, where essential traits to interact with Legumes are plasmid-encoded. Symbiosis plasmids are likely to evolve into stably inherited domesticated plasmids (termed chromids). Recent research reveals that potential chromids are widely distributed among plant-associated bacteria. Studying the origin of plasmids in the plant-associated genus *Pantoea*, we show that the large *Pantoea* plasmid (LPP) and plasmid pPag1 are domesticated. Phylogenomic analyses trace back the plasmid origin to ancestral nodes in the *Pantoea* species tree. We reconstruct main events in chromid evolution, including massive gene gain, some redundant with the chromosome, and differential gene loss.

## Introduction

Plasmids are extrachromosomal genetic elements that reside in prokaryotic organisms, where they replicate using the host replication machinery. Plasmid persistence within the host depends on their ability to replicate and segregate into daughter cells during cell division (1). Alternatively, plasmids can confer their host a fitness advantage over non-hosts in the population via addiction mechanisms (2), or the expression of traits beneficial to the host (e.g., lactic acid metabolism in *Bacillus lactis* (3)). Plasmid-host adaptation over long time scales may lead to the evolution of domesticated plasmids that are an integral component of the host genome in the form of chromids (4, 5). The coordination of plasmid replication with cell division, which ensures faithful plasmid segregation, is a crucial step in the emergence of domesticated plasmids. In *Vibrio cholera*e, chromid replication is well coordinated with the chromosome replication via a check-point mechanism that is encoded in the chromosome (6). The evolution of plasmid domestication may be furthermore accompanied by non-functionalization and loss of plasmid conjugation genes (7, 8). Domesticated plasmids typically confer their host traits that are essential in the host’s natural habitat. Examples are nodulation and nitrogen fixation in Rhizobia (9, 10), virulence factors and biofilm formation in enterohemorrhagic Escherichia (11) and chitin utilization in *Vibrio cholera* (12), where the plasmid acquisition likely facilitated the host habitat expansion.

A main characteristic of chromids is the presence of core genes that are universally present in all taxon members (4, 5). The evolutionary origin of core genes in chromids has been debated in the literature where two competing hypotheses have been suggested. According to the ‘translocation hypothesis’ essential core genes in chromids result from the translocation of chromosomal genes to the chromid following plasmid domestication (5). Alternatively, the ‘redundancy hypothesis’ posits that the evolution of core genes in chromids entails preliminary genetic redundancy between the plasmid and chromosome that is followed by differential loss (5). According to the redundancy hypothesis, precursors of domesticated plasmids are megaplasmids that harbor homologs to chromosomal genes; if the chromosomal and plasmid genes are functionally redundant, a loss of the chromosomal homolog is expected to have a negligible effect on the host phenotype. A gradual process of redundant chromosomal gene loss is thus expected to promote the evolution of plasmid indispensability to the host. Nonetheless, evidence in support of either hypothesis remains scarce.

Members of the genus *Pantoea* are widely associated with plants (13). Several *Pantoea* species are recognized plant pathogens, including *P. anantis*, that infects mono-and dicotyledonous plants (14, 15), the maize pathogen *P. stewartii* (16) and the onion pathogen *P. allii* (17). In contrast, strains among the species *P. vagans, P. agglomerans, P. eucalypti, P. eucrina* and *P. dispersa* are generally recognized as commensals (18–22), yet pathogenic *P. agglomerans* have been reported in these species (23). Plant growth promoting *Pantoea* harbor traits such as nitrogen fixation (22, 24), phosphate solubilization (20), production of the plant hormone indole-3-acetic acid (25), heavy metal tolerance (21), antifungal activity (26, 27) and antibiotics activity, e.g., against the causative agent of fire blight in trees (28). Many *Pantoea* species harbor a megaplasmid named ‘Large *Pantoea* Plasmid’ (LPP) and a medium sized plasmid named pPag1 (29). Homologs of plasmid pPag1 (termed also LPP-2) are widely present in *P. agglomerans*; besides their putative contribution to sucrose utilization, not much is known about traits conferred by pPag1 (30). In contrast, curing of LPP has a significant effect on *Pantoea* physiology. LPP curing in the maize pathogen *P. ananatis* strain DZ-12 (termed pDZ-12), leads to a decrease in biofilm formation and decreased disease symptoms in infected maize plants (31). Similarly, curing of LPP in *P. vagans* strain C9-1 led to reduced (or loss of) ability to utilize several carbohydrate sources, e.g., maltose and salicin (32), reduced swarming motility, and reduced Indole production (33). The LPP thus harbors multiple traits that are relevant for the *Pantoea* endophytic lifestyle. The wide distribution of plasmids LPP and pPag1 in the phylum and their conservation across *Pantoea* species suggests that they have been domesticated during the evolutionary history of the genus (29). Here we reconstruct the evolution of LPP and pPag1 based on comparative genomics of nine *Pantoea* species. Analyzing the phylogenies of plasmid genes, we examine evidence in support of alternative hypotheses for the emergence of domesticated plasmids.

## Results

### The acquisition and diversification of plasmids LPP and pPag1 coincide with *Pantoea* speciation events

To investigate the evolution of *Pantoea* plasmids, we clustered 462 plasmids found in 253 *Pantoea* isolates into 39 clusters of homologous plasmids according to their gene content (Table S1). Homologs of LPP and pPag1 plasmids correspond to 80% of the *Pantoea* plasmids and are characterized by similar gene content (Fig. 1a). LPP homologs (214 plasmids) were clustered into six distinct groups, where all plasmids harbor a replication initiation protein (RepB) that is homologous to previously annotated LPP plasmids (29). The clustering of 114 pPag1 homologs yielded two distinct groups. All pPag1 plasmids harbor a homolog to the previously annotated pPag1 RepB. The remaining 134 plasmids were clustered into 31 small groups having low connectivity with the other groups (Fig. 1a).

**Fig 1.**
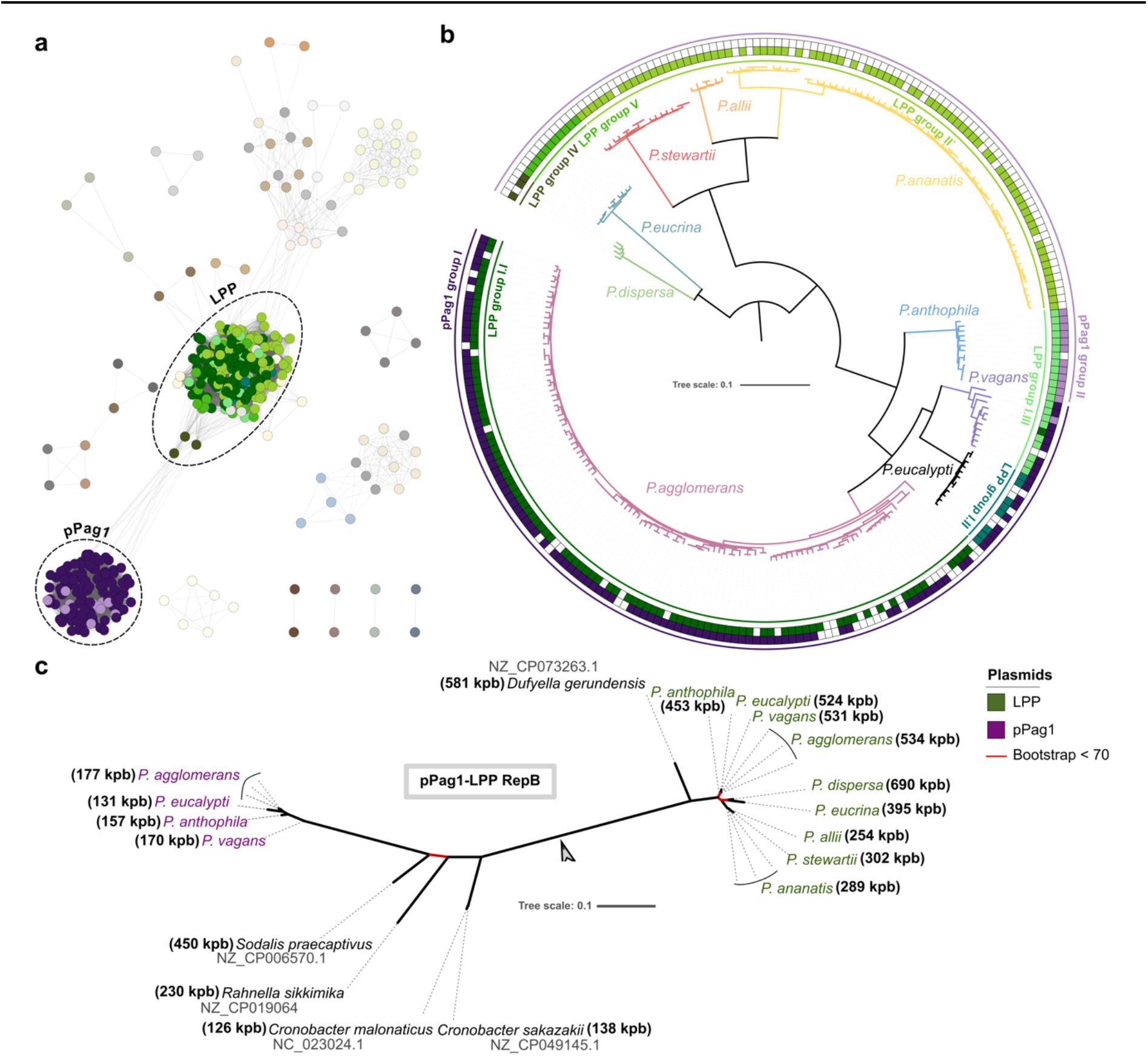
The LPP and pPag1 groups correspond to *Pantoea* species taxonomy. (**a**) A network of plasmid gene content similarity. Nodes in the network correspond to plasmids and the edges correspond shared gene content calculated as the overlap coefficient (see methods). Node color is according to the LPP and pPag1 taxonomic units shown in (b). Edge length is proportional to plasmid similarity, such that similar plasmids appear as aggregates in the network. Edge thickness is proportional to the overlap coefficient value. Only edges having overlap coefficient ≥ 0.4 are shown. (**b**) Phylogenetic tree of 253 *Pantoea* strains from nine species isolated mostly from plant hosts (Table S1). The tree was inferred from 299 chromosomal single copy gene families. The root was inferred using a phylogenomic rooting approach and according to a recently published phylogeny of the genus (see Supplementary Note 1 (34, 35)). The tree outer rings show the distribution of LPP (in green) and pPag1 (in purple) groups. (**c**) Phylogeny of *repB* homologs in LPP and pPag1 of 13 representative *Pantoea* species (Table S1). LPP homologs are shown in green labels; pPag1 homologs are shown with purple labels. The root was inferred using MAD (36). The plasmid size is shown for all species. For *Pantoea* species, plasmid size corresponds to the median plasmid size per species. Branches highlighted in red have bootstrap values under 70% (using 1000 replicates).

To reconstruct the LPP and pPag1 evolutionary history, we first inferred the *Pantoea* species phylogeny. The species tree topology comprises clades that correspond to the nine *Pantoea* species (Fig. 1b). The inferred root of the species tree splits *P. eucrina* and *P. dispersa* from the remaining species (Fig. 1b; Supplementary Note 1; Fig. S1; Table S2). The distribution of LPP and pPag1 clusters across the *Pantoea* species tree suggests the co-divergence of both plasmids with the *Pantoea* species (Fig. 1b). The prevalence of LPP adds support to the view that the LPP origin trace back to the genus origin (29). Here we term the LPP clusters based on previous classification of LPP groups (29). LPP plasmids in group I were clustered into three clusters that are supported by the RepB phylogeny (Fig. S2). Plasmids in LPP group I.I are specific to *P. agglomerans*, group I.II plasmids were observed in *P. eucalypti*, and group I.III plasmids are shared to *P. vagans* and *P. anthophila*. Plasmids in group II were observed only among *P. ananatis, P. allii* and *P. stewartii*. Our analysis uncovers two additional LPP groups: group IV in *P. dispersa* and group V that is specific to *P. eucrina*. In addition to gene content differences, the LPP clusters differ in their genome size (Fig. S3), where group II in *P. ananatis, P. stewartiii* and *P. allii* were smaller compared to group IV in *P. dispersa*. Homologs of pPag1 plasmid were observed only in *Pantoea* isolates from among *P. agglomerans, P. vagans, P. anthophila* and *P. eucalypti*. The RepB phylogeny support the presence of two pPag1 groups (Fig. S2). pPag1-group I is shared among *P. agglomerans, P. vagans* and *P. eucalypti*, while pPag1 group II is specific to *P. anthophila* isolates. The distribution pPag1 homologs within the *Pantoea* species suggests that the pPag1 acquisition corresponds to the divergence of pPag1 hosts within the genus. Furthermore, the LPP and pPag1 clusters are largely matching main divergence events in the *Pantoea* species phylogeny, suggesting that both plasmids are vertically inherited.

What is the origin of LPP and pPag1? The replication initiation proteins of LPP and pPag1 are members of the same RepB-type family and exhibit considerable sequence similarity (e.g., 66% identical amino acids over 90% of the LPP *repB* length with the pPag1 *repB* in *P. agglomerans* R1). Thus, both plasmids are classified into the same plasmid incompatibility group, IncFIB, suggesting a common plasmid origin. To examine the evolutionary relations between LPP and pPag1, we inferred a joined RepB phylogeny including additional homologs that were identified outside of the *Pantoea* genus. The sequence similarity search against plasmid sequences yielded only few putative homologs. The LPP *repB* genes had homologs in the megaplasmid of the plant-growth promoting *Duffyella gerundensis*, a sister species of *Pantoea*, within the same *Erwiniaceae* family (37, 38). Two homologs of LPP *repB* were identified in *Cronobacter* plasmids that were previously classified as IncFIB incompatibility group (39). Putative homologs of the pPag1 *repB* were identified in a megaplasmids from *Sodalis praecaptivus*, which was isolated from a hand wound following impalement with a tree branch and is closely related to *sodalis*-allied insect endosymbionts (40, 41). Another homolog was identified in *Rahnella sikkimica*, which was isolated from soil yet its gene content hints a plant-associated lifestyle (42). The root of the *repB* phylogeny was inferred at the split between the LPP and pPag1 homologs (Fig. 1c), indicating an independent origin of the two plasmids in *Pantoea*. Considering the rare homologs to the LPP and pPag1 RepB that we identified here, it is tenable to hypothesize that the LPP and pPag1 lineages originate in plasmids that reside in plant-associated bacteria.

### LPP and pPag1 are vertically inherited within their *Pantoea* host lineage

To examine evidence for vertical inheritance in the evolution of LPP and pPag1, we compared their phylogeny with that of the *Pantoea* chromosome. The phylogenies of the three replicons (i.e., both plasmids and the chromosome) are largely compatible, except for two splits (marked (3) and (8) in Fig. 2a-c). To quantify the incongruence among the three replicon phylogenies, we examined the individual phylogenies of complete single-copy (CSC) gene families. The split of *P. eucalypti* and *P. vagans* (marked (3)) was observed in the chromosome and LPP phylogenies but not in the pPag1 phylogeny. That split is well supported by chromosomal and LPP CSC gene trees and had a lower support in the pPag1 CSC gene trees (Fig. 2d). The other incompatible split corresponds to the split of *P. anthophila* and *P. vagans* from the remaining species in the pPag1 phylogeny (marked (8)), that is not present in the chromosome or the LPP phylogenies. This split is supported by about half of the CSC genes in pPag1, and can be observed in about a third of the chromosomal and LPP CSC gene trees (Fig. 2d). Taken together, the inferred split between *P. eucalypti, P. vagans* and *P. anthophila* and the remaining species is well supported by the data, while conflicting splits in gene trees show evidence for an ongoing process of incomplete lineage sorting among these three species.

**Fig 2.**
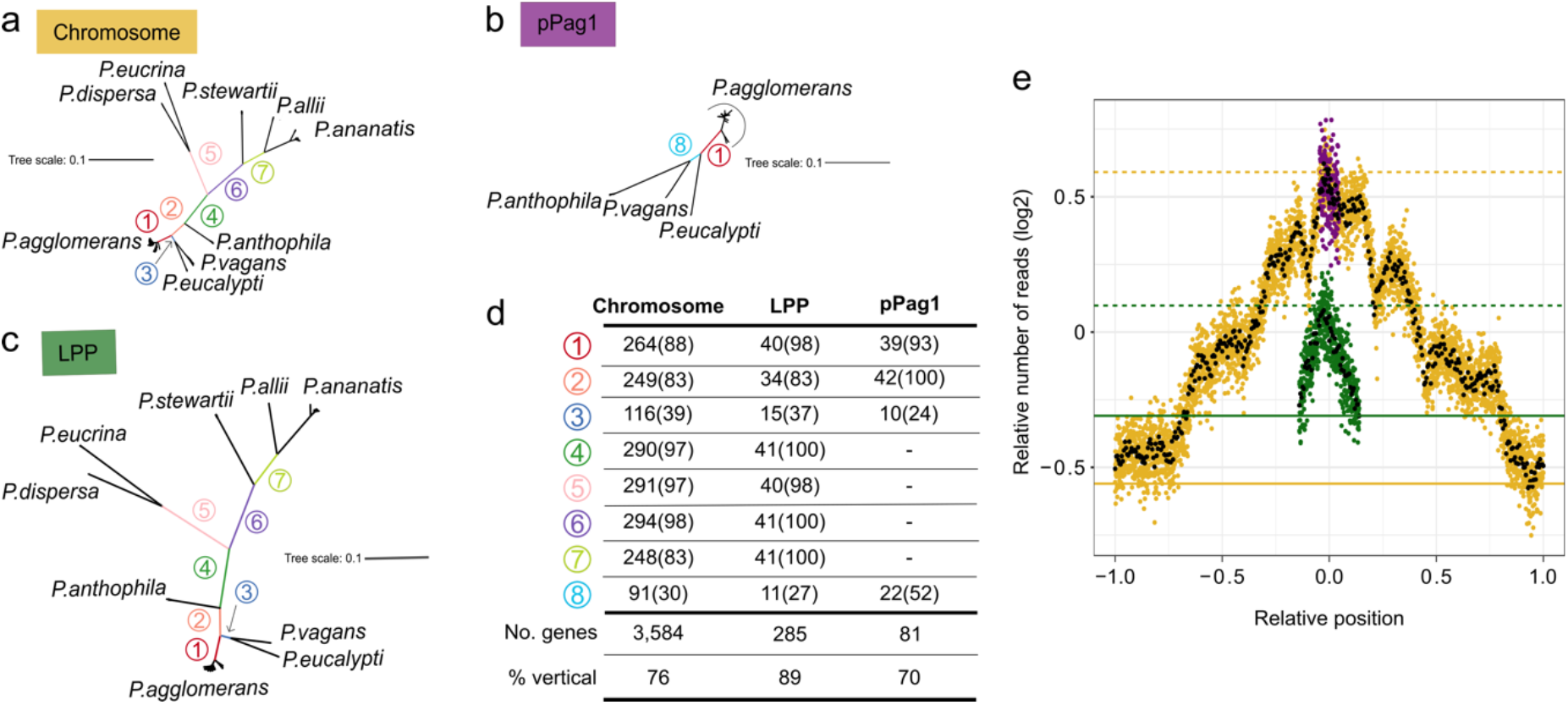
LPP and pPag1 are vertically inherited plasmids. Replicon phylogenies were inferred from complete single-copy (CSC) gene families that are exclusively present in the focal replicon. Internal branches that correspond to major species divergence events are highlighted. (**a**) Chromosome phylogeny reconstructed from 299 CSC gene families present in 253 isolates. (**b**) LPP phylogeny reconstructed from 41 CSC gene families in 143 isolates. (**c**) pPag1 phylogeny reconstructed from 42 CSC gene families present in 87 isolates. The number of isolates used for the analysis of each replicon was adjusted in order to increase the number of CSC families. (**d**) Number (and percentages) of CSC gene trees where the split was observed. The bottom lines show the total number (and percentage) of gene trees having a topology not significantly different from the species tree topology (i.e., their evolution is dominated by vertical inheritance). (**e**) Marker frequency analysis (MFA) for LPP (green), pPag1 (purple) and chromosome (yellow). The number of reads is presented for 1 kb (colored dots) and 10 kb (black dots) windows as the log_2_ ratio of the number of reads divided by the number of reads of a non-replicating sample. Additionally, the number of reads is normalized by the total number of reads per sample. The x-axis indicates the relative position on the chromosome, where 0 corresponds to the origin of replication in all three replicons, and 1 and -1 correspond to the replication termination in the chromosome. Dashed lines indicate the origin of replication while continuous lines indicate termination in LPP and pPag1.

To examine further evidence for co-divergence of LPP, pPag1 and their hosts, we tested for compatibility of plasmid single-copy gene trees with the main speciation events as observed in the chromosome phylogeny. The test was applied to replicon-specific single-copy gene families (i.e., complete and partial) comprising ≥4 sequences from at least two species. Constraining the tree topologies according to the species topology showed that the topology of most single-copy gene families is not significantly different from the species phylogeny (Fig. 2d). The LPP and pPag1 phylogenies indicate the presence of mechanisms for stable plasmid inheritance of both plasmids.

To test if the plasmid replication is coordinated with the chromosome replication, we examined the replication pattern of the two plasmids and the chromosome in the wildtype *P. agglomerans* strain R1 during early exponential growth using a marker frequency analysis (MFA (6, 43)). The results from two independent replicates show that LPP replication was initiated after 43% or 47% of the chromosome have been replicated (Fig. 2e; Fig. S4). The similarity among the replicates indicates that the replication of LPP is synchronized with the chromosome. The replication pattern of pPag1 appears highly similar between the two MFA replicates, however the pPag1 small size does not enable a reliable evaluation of synchronized replication with the chromosome. Taken together, we conclude that the evolution of both plasmids is governed by vertical inheritance, a key hallmark of plasmid domestication.

### The LPP pangenome is shared with the chromosome and comprises core genes

To evaluate the plasmid contribution to the *Pantoea* pangenome, we examined the distribution of gene families in plasmids and chromosome. About 10% of the gene families include homologs in both the chromosome and the LPP (Fig. 3a; Fig. S5a). The core *Pantoea* genome at the genus level includes 191 gene families shared between the chromosome and the LPP, where 20 gene families comprise only LPP genes. The number of core gene families (present in ≥95% isolates) shared between the chromosome and LPP is lower at the species level and ranges between 25 gene families in *P. eucrina* and 94 gene families in *P. ananatis*. In contrast, the number of core gene families found exclusively in the LPP is higher at the species level, ranging between 37 gene families in *P. agglomerans* and 446 gene families in *P. dispersa* (Fig. 3a). The distribution of shell gene families (present in 10-95% isolates) is likewise characterized by high frequency of LPP gene families at the species level, while the cloud gene families (present in ≤10% isolates) are mostly replicon specific (Fig. S5a). Gene families shared exclusively between pPag1 and the chromosome are rather rare in the *Pantoea* core genome (21 families; Fig. 3a). Nonetheless, gene families including exclusively pPag1 genes are found in the core genome of *P. vagans* and the shell genome of *P. vagans, P. agglomerans, P. eucalypti* and *P. anthophila*. Gene families shared exclusively between the LPP and pPag1 are found only in the shell and cloud pangenomes (51 gene families). Taken together, the *Pantoea* pangenome is distributed across LPP, pPag1 and the chromosome, where plasmid-specific gene families are observed depending on the taxonomic level. The presence of pPag1-specific genes in the pPag1 host species adds support to our inference of pPag1 vertical inheritance, while the presence of LPP-specific families at the genus level is in accordance with ancient origins of the LPP.

**Fig 3.**
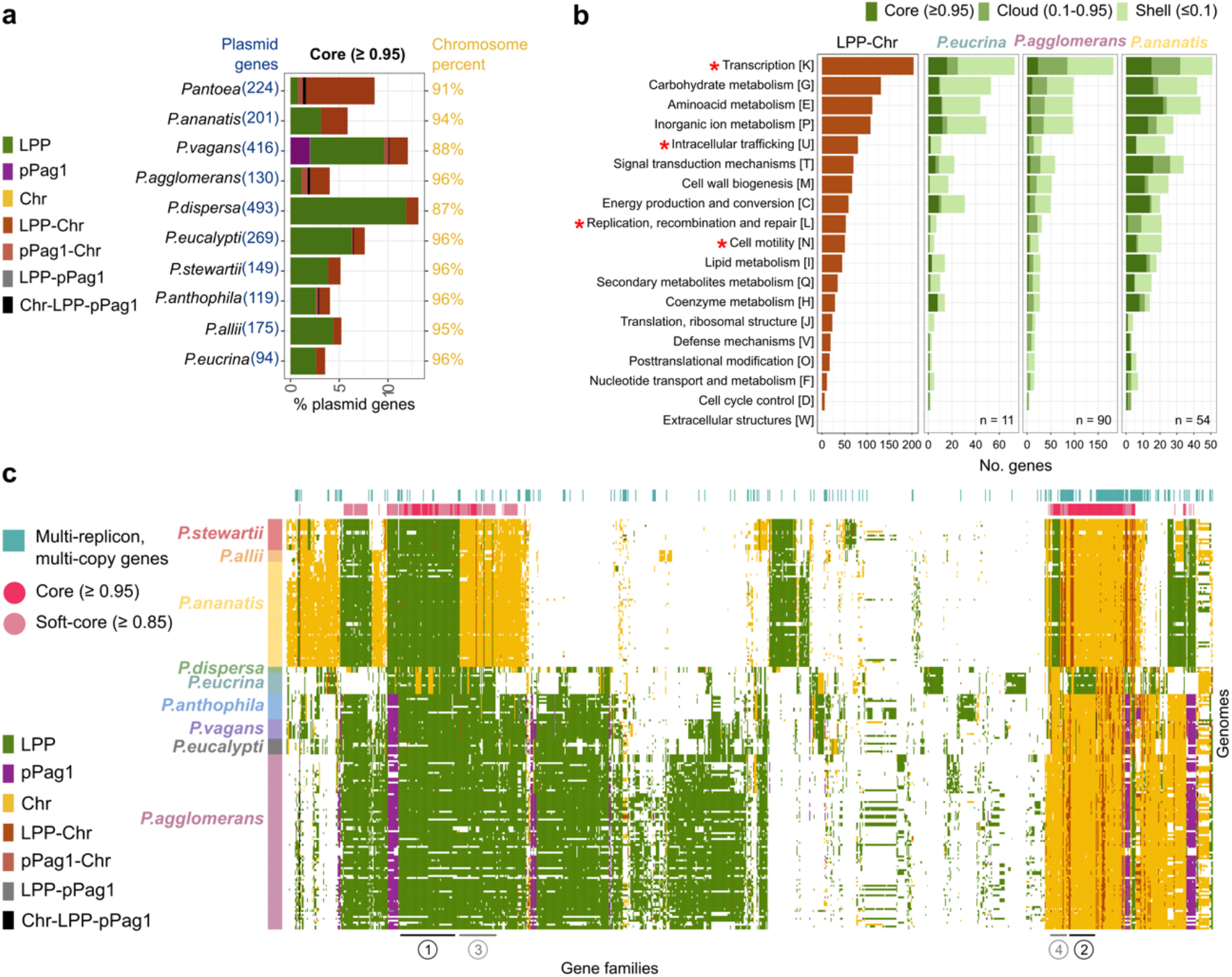
The LPP pangenome is shared with the chromosome and comprises a high frequency of transcription-related genes. (**a**) Distribution of core genes in the *Pantoea* pangenome in 211 isolates harboring the LPP plasmid. Three out of the 214 original isolates carrying the LPP were excluded from this analysis due to potential assembly artifacts (Fig. S6 and Fig. S7). The stacked bar graph shows the proportion of gene families according to replicons where family-members are observed. (**b**) Functional annotation of genes shared between LPP and chromosome and the LPP pangenome in representative species: *P. eucrina, P. agglomerans* and *P. ananatis*. The number of isolates (n) in the analysis is shown below. COG categories where the frequency of genes in that category is significantly different among all nine species are marked by a red asterisk. (**c**) Replicon presence/absence pattern (PAP) of gene families in the LPP pangenome. Rows correspond to isolates and columns correspond to gene families. The data presented correspond to 1,086 protein families including an LPP homologs in at least 10 isolates. The rows are ordered according to the species phylogeny in (Fig. 1b). Clusters of gene families characterized by species-specific gene location in plasmids and chromosome are indicated by numbers below the heatmap.

The functional composition of gene families shared between LPP and the chromosome is similar to the functional composition of the LPP pangenome at the species level (Fig. 3b, Fig. S5b). Most gene families shared between LPP and the chromosome are annotated to have transcription-related functions, followed in frequency by gene families classified as carbohydrate and amino-acid metabolism (Fig. 3b). Additionally, transcription-related functions are prevalent in multicopy gene families that harbor homologs in the LPP and the chromosome of the same isolate. The proportion of LPP gene families classified into the functional categories is similar among *Pantoea* species for most categories (Fig. 3b; Fig. S5b; Table S3). The distribution of transcription-related gene families stands out as an exception: in *P. ananatis, P. allii*, and *P. stewartii* the frequency of chromosomal gene families in the transcription category is markedly higher compared to LPP while in the remaining species the frequency of LPP gene families in the transcription is slightly higher than the chromosomal ones. The differences in LPP functional composition among *Pantoea* species are in line with species-specific evolution of the LPP taxonomic units (Fig. 1b).

To further examine the evolution of LPP gene families, we compared the genomic location of all gene families that include at least ten LPP-encoded genes among *Pantoea* species. The replicon presence/absence pattern (PAP) reveals core *Pantoea* gene families comprising LPP homologs that also include chromosomal and pPag1 genes. The PAPs reveal two main patterns of gene families comprising LPP and chromosomal homologs, which are reminiscent of the main *Pantoea* species phylogeny (Fig. 3c). The first PAP is characterized by a combination of chromosomal homologs in *P. dispersa* and *P. eucrina* and LPP homologs in the remaining species, or the other way around: LPP homologs in *P. dispersa* and *P. eucrina* and chromosomal homologs in the remaining species (groups 1 and 2 in Fig. 3c); this PAP is matching to the basal *Pantoea* phylogeny split (Fig. 1b). The second pattern corresponds to core genes that are located either on the chromosome in *P. ananatis, P. allii* and *P. stewartii* and the LPP of the remaining species or alternatively, genes that are located on the LPP of *P. ananatis, P. allii* and *P. stewartii* and the chromosome of the remaining species (groups 3 and 4 in Fig. 3c). That pattern matches the split between *P. ananatis, P. allii* and *P. stewartii* and the remaining species in the species phylogeny. The correspondence between the replicon PAPs and main splits in the *Pantoea* phylogeny suggests that gene family diversification coincides with two main speciation events in the genus. Notably, most of the gene families comprise a homolog on a single replicon within each isolate, with a total of 465 (14%) gene families including homologs on both LPP and the chromosome within the same isolate (see multi-replicon, multicopy gene families in Fig. 3c). Hence, the presence of duplicated genes in multiple replicons within *Pantoea* genomes is rather rare.

Taken together, the *Pantoea* pangenome comprises core LPP genes at the genus level that likely correspond to LPP-mediated novel gene acquisition in the common *Pantoea* ancestor. Gene families comprising chromosomal and LPP homologs are the result of either gene translocation to, or from, LPP, or alternatively differential loss of genes that were redundant in the chromosome and LPP. The phyletic pattern observed in the replicon PAPs indicates that LPP gene acquisition, translocation and loss coincided with species divergence events in *Pantoea*.

### Phylogenies of LPP genes provide support for the genetic redundancy hypothesis

To test if the LPP evolution is better described by the ‘translocation hypothesis’ or the ‘redundancy hypothesis’, we examined evidence for gene translocation and differential loss in the evolution of shared core LPP and chromosome gene families. To constrain the effect of phylogenetic biases in the analysis (e.g., due to a large number of OTUs (44)), we analyzed a reduced dataset of 13 genomes of representative *Pantoea* species (Table S1). The largest category of gene families has a replicon PAP that is matching the basal *Pantoea* phylogeny split (40% complete single copy genes and 43% partial single copy genes; Fig. 4a). The second most frequent gene families category has a replicon PAP that is matching the split between *P. ananatis, P. allii* and *P. stewarti* and the remaining species (44% complete single copy genes and 22.6% partial single copy genes; Fig. 4a). Hence, the main replicon-specific PAPs of gene families observed in the preliminary dataset are prominent also in the reduced dataset.

**Fig 4.**
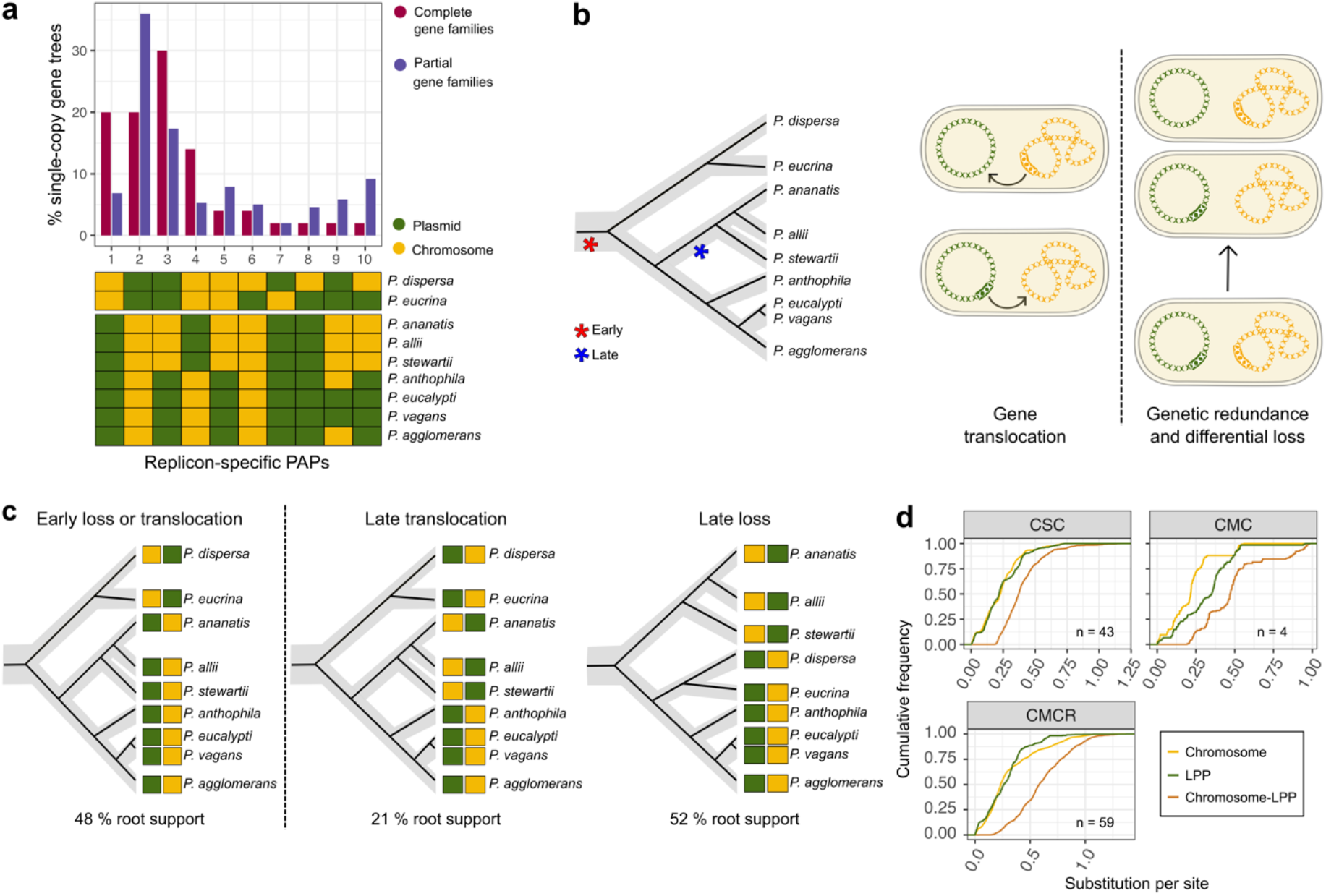
Phylogenetic analysis and genetic distances provide support for the redundancy hypothesis in LPP evolution. **(a)** Distribution of single copy gene families in the LPP pangenome that comprise chromosomal homologs. Highly frequent replicon-specific PAPs were identified in 50 complete single copy gene families that are present in all nine species. A total of 183 partial single copy gene families found in at least 4 species, were mapped onto the replicon-specific PAPs. The replicon-specific PAPs are displayed in a heatmap form. **(b)** An illustration of the tested evolutionary hypotheses. Putative branches matching the occurrence of early and late events matching the most common PAPs found among single-copy gene families are marked by an asterisk. The cartoons illustrate hypotheses of gene translocation between the LPP and chromosome (left) and ancestral genetic redundancy followed by differential loss (right). **(c)** Expected gene tree rooted topologies and PAPs for the tested scenarios (illustrated in (**b**)). The percentage of genes in (**a**) that support each root position is reported below (see detailed report in Table S4). **(d)** Distribution of pairwise phylogenetic distance calculated as substitution per site across complete gene families shared between chromosome and LPP plasmid. Distance distributions are presented for complete single-copy gene families (CSC); complete multicopy gene families, where copies are present in a single replicon (CMC) and complete multicopy gene families, where copies can be present in both replicons (CMCR). The number of gene families analyzed (n) is presented.

In accordance with the most frequent patterns of gene family distribution in LPP and the chromosome, we hypothesized two alternative evolutionary scenarios. The *early translocation or loss* scenario is proposed for gene families where the replicon-specificity corresponds to the basal species phylogeny split (PAPs 1, 2 in Fig. 4a and groups 1, 2 Fig. 3c). According to this *early* scenario, the LPP acquisition was accompanied either by gene translocation between LPP and the chromosome, or, if a chromosomal homolog existed, the LPP acquisition was followed by loss of either the chromosomal or the plasmid homolog (Fig. 4b). The second scenario -*late translocation or loss* - is proposed for gene families where the replicon-specificity matches the split between *P. ananatis, P. allii* and *P. stewartii* and the remaining species (PAPs 3, 4 in Fig. 4a and groups 3, 4 in Fig. 3c). According to the *late* scenario, the translocation or loss event occurred during the divergence of *P. ananatis, P. allii* and *P. stewartii* ancestor (Fig. 4b). Note that the most frequent PAP points towards the loss or translocation of plasmid genes rather than the loss or translocation of chromosomal genes in this lineage (see PAP 3 vs. PAP 4 Fig. 4a), which matches the observation of smaller LPP size in *P. ananatis, P. allii* and *P. stewartii* (Fig. S3). The phylogenies of gene families having a PAP matching to the *early translocation or loss* scenario are expected to have a similar topology to that of the species tree, that is, with a root position matching the basal species phylogeny split. Indeed, we find that the most frequently observed root position in these gene families is the basal *Pantoea* split (48% of gene trees), followed by a root position on the branch leading to *P. ananatis, P. allii* and *P. stewartii* (22% gene trees) (Fig. 4c; Table S4). Furthermore, the basal *Pantoea* split is significantly supported as the root position among phylogenetic trees reconstructed from the *early* PAPs (FDR adjusted P-value = 0.0221, using two-sided Wilcoxon test). Note that the *early* tree topology (Fig. 4c) does not enable distinguishing between translocation or loss events. In contrast, the root inference for gene families matching the *late* scenario enables a distinction between translocation and loss, since these two events induce different root positions (Fig. 4c). The most frequent root position in the *late* gene families is the branch that splits chromosomal and plasmid homologs (52% of the gene trees), followed by a root position on the branch leading to *P. eucrina* and *P. dispersa* (21% gene trees; Fig. 4c; Table S4). Furthermore, the root position expected for *late loss* is significantly more supported in trees of the *late* PAP category compared to alternative root positions (FDR adjusted P-value = 0.0218, using two-sided Wilcoxon rank sum test). Thus, we conclude that gene loss, rather than gene translocation, was the most frequent event in the evolution of the *late* gene families.

The most frequent rooted tree topologies in both *early* and *late* scenarios comprise a deep split between chromosomal and plasmid homologs, which appears in most gene trees having more than one plasmid homolog (68%; 186 genes). To further distinguish between scenarios of gene loss or translocation, we compared the pairwise phylogenetic distance within and between homologs present in the chromosome and LPP (Fig. 4d). The distance between homologous gene pairs in the same replicon in complete single copy families was significantly smaller than the distance between gene pairs in different replicons (P-value < 2.2×10^−6^, using two-sided Wilcoxon test). A significant difference was furthermore observed in all gene family categories, including partial single-copy and multicopy gene families, as well as those gene families that harbor chromosomal and LPP homologs within the same strain (Fig. 4d; Table S5; Fig. S8). Comparing the distance distribution to chromosome-specific gene families further shows that the observed differences in phylogenetic distances between chromosomal and LPP genes are significantly larger than the expected distances by species divergence alone (P-value < 2.2×10^−6^, using two-sided Wilcoxon test; Fig. S9). These results indicate that the divergence of chromosomal and LPP genes preceded the *Pantoea* species divergence events.

Taken together, we conclude that the LPP diversification occurred during two main evolutionary events matching early and late species divergence episodes. The results of the phylogenetic analysis and comparison of gene distances are in support of early divergence of chromosomal and LPP genes and differential loss of homologs following the LPP acquisition, as expected according to the redundancy hypothesis for chromid evolution.

### The pPag1 evolution reveals early stages of plasmid domestication *en route* to a chromid

The plasmid pPag1 has a limited distribution within the genus and its contribution to *Pantoea* physiology remains understudied. Nonetheless, its presence in species sharing a common ancestor and our inference of pPag1 vertical inheritance suggests that pPag1 plasmid is domesticated in the analyzed host species (Fig. 1b; Fig. 2). The functional composition of pPag1 is similar among *Pantoea* species (Fig. S10), which in support of a recent acquisition (compared to LPP). The majority of pPag1 gene families shared with the chromosome encode for transcription-related protein functions, followed by intracellular trafficking, cell motility and signal transduction mechanisms (Fig. S10). Examining the replicon PAP of pPag1 gene families reveals four main PAP patterns. The most frequent PAP corresponds to gene families that are mostly pPag1-specific (Fig. 5a). Two additional frequent PAPs correspond to gene families comprising pPag1 and chromosomal homologs; one of these PAPs contains several gene families that have duplicates in the same isolates (group 1, Fig. 5a). A total of 113 (25%) gene families in the pPag1 pangenome have chromosomal homologs in the same isolate (see multi-replicon, multi-copy genes in Fig. 5a). Thus, gene redundancy with the chromosome is higher in the pPag1 pangenome compared to the LPP pangenome. The second PAP comprises gene families that have a chromosomal homolog in the genomes of those species that do not harbor pPag1 (group 1, Fig. 5a). Similarly to LPP, the phylogenetic distance between homologous genes in the same replicon (chromosome or pPag1) is significantly smaller compared to the distance between genes in different replicons (P-value < 2.2×10^−6^, using two-sided Wilcoxon test; Fig. 5b). This difference indicates that pPag1 and chromosomal homologs diverged prior to the diversification of pPag1-hosts within the *Pantoea* genus. Thus, we conclude that the pPag1 acquisition introduced genetic redundancy that was accompanied by differential loss of homologous genes in the chromosome and pPag1, as expected according to the redundancy hypothesis.

**Fig 5.**
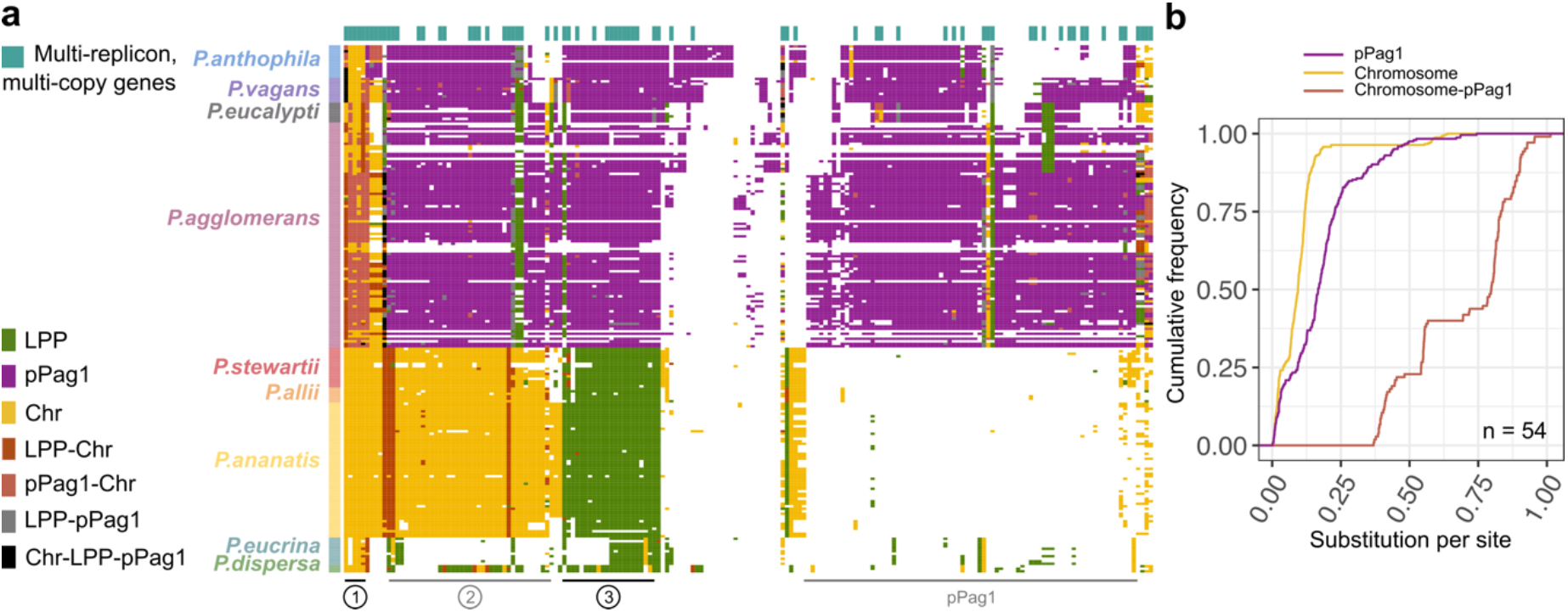
The pPag1 pangenome shows a signature of early stages of plasmid domestication. **(a)** Presence / absence pattern of pPag1 pangenome according to replicons. The analysis was restricted to 211 isolates included in **Fig 3c** and 189 protein families which are present in at least 10 pPag1 plasmids. Each row corresponds to a genome and each column corresponds to one gene family. The rows are clustered according to the species phylogeny (**Fig. 1b**). Groups of gene families characterized by species-specific replicon PAP are indicated by numbers below the heatmap. **b**, Distribution of pairwise phylogenetic distance calculated as substitution per site across gene families shared between pPag1 and the chromosome. Only species carrying the pPag1 plasmid were considered in the analysis.

The replicon PAP reveals a fourth PAP pattern of 51 gene families shared between pPag1 and the LPP of pPag1-less species (group 3, Fig. 5a). These gene families include several typical plasmid functions (e.g., active partition and toxin-antitoxin systems). Plasmid pPag1 furthermore harbours several antibiotic resistance genes, e.g., homologs of Polymyxin resistance genes (*arnBCADTEF;* (45)). Moreover, a homolog of organophosphorus hydrolase (*opdE*) is widely present in the pPag1; a homolog of this enzyme from the plant associated *Leuconostoc mesenteroides* was recently shown to degrade diverse insecticides (46).

The similarity in the pattern of pPag1 gene sharing with the chromosome to that of LPP and the evidence for pPag1 vertical inheritance (Fig. 2) indicate that the pPag1 is a domesticated plasmid. Compared to LPP, the pPag1 acquisition was late in the *Pantoea* evolution; the pPag1 contribution to the *Pantoea* species core genome is rather small and pPag1 gene redundancy (with the chromosome) is more frequent compared to LPP. Taken together, the pPag1 pangenome properties support the inference that pPag1 is at an early stage of plasmid domestication, *en route* to becoming a chromid.

## Discussion

The evolution of adaptive traits via horizontal gene transfer can facilitate the acquisition of novel ecological niches in prokaryotes (47, 48). Horizontal gene transfer is a major driver of prokaryote speciation, however, most models of species divergence focused on the role of homologous recombination (49–52), while plasmid acquisition was considered as a driver of species divergence in only few taxa, e.g., Rhizobia (53) and Yersinia (54, 55). The *Pantoea* species we examined here are frequently associated with plants. Nonetheless, species in the genus are known to be found in diverse habitats (19). A recent phylogenetic reconstruction of the genus evolutionary history shows species not associated with plants in a basal position, thus pointing towards *Pantoea* adaptation to the plant habitat as a later adaptation. Our results hints at a contribution of LPP and pPag1 to the adaptation of a plant-associated lifestyle in *Pantoea*.

The presence of plasmid genes in the species core genome is a hallmark of bacterial taxa that harbor chromids (4) (or domesticated plasmids). Core gene families comprising plasmid genes are of two types: gene families that are exclusively present in the plasmid and gene families comprising homologs in both the plasmid and chromosome. The latter type of mosaic plasmid-chromosome gene families makes up the majority of the core plasmid genes in *Pantoea* (Fig. 3). The emergence of plasmid-specific core genes in the species pangenome is well explained by species divergence following plasmid acquisition. But what is the origin of mosaic plasmid-chromosome core gene families? The translocation hypothesis posits the occurrence of gene transfer or translocation from the chromosome to the plasmid (56). However, a recent study suggests that gene transfer between plasmids and chromosomes is rather rare and mostly depends on gene association with mobile genetic elements (e.g., transposons) (57). Hence, while translocation of genes from chromosome to the chromid has been reported in Rhizobia and *Burkholderia* (58, 59), such instances may be anecdotal. Our inference for the evolution of mosaic plasmid-chromosome core gene families is rather in agreement with the redundancy hypothesis that implicates genetic redundancy and differential loss (60). The redundancy hypothesis is further supported by the observation that ca. 14% of the chromosomal genes in *S. meliloti* are functionally redundant with the resident plasmids (61). Notably, in most of the mosaic core gene families we identified a robust split between the plasmid and chromosome homologs, that is highly supported by phylogenetic inference and comparison of genetic distances (Fig. 4d). Our results thus indicate that chromosomal and plasmid homologs in *Pantoea* diverged prior to main speciation events in the genus. The plasmid homologs in those core gene families shared between plasmid and chromosome are thus better described as xenologs (62), rather than paralogs.

The evolution of genetic redundancy has been extensively studied in the context of whole genome duplication (WGD) in eukaryotes. Example are studies of WGD in the evolution of yeast (63) and Arabidopsis (64), which showed that most duplicated genes are differentially lost over time and return to a single copy state (65). An important difference between WGD and megaplasmid acquisition is the level of genetic redundancy, which is a determinant of the fate of duplicated genes (66). WGD-derived paralogs are preliminarily identical while chromosomal genes and the plasmid xenologs are most likely preliminarily diverged, both in their sequence (Fig. 4d and Fig. 5b) as well as the mode of transcriptional regulation. Our inference points towards differential loss of duplicated genes following the LPP (or pPag1) acquisition, similarly to post-WGD event. The *late* scenario we tested here (Fig. 4) coincides with the emergence of commonly recognized plant pathogens in the species *P. ananatis, P. allii, and P. stewarti*. It is tenable to hypothesize that the LPP size reduction in those species is associated an adaptation of a parasitic lifestyle in those species. Our results further show a high frequency of redundancy in gene families involved in transcription-related functions. These findings are in agreement with previous reports on the enrichment of transcriptional regulators in chromids (5). The retention of functionally redundant duplicated genes potentiates the evolution of transcriptional plasticity that can contribute to rapid physiological response to diverse environmental conditions (67). Phenotypic plasticity is an important property in bacteria leading a bi-phasic lifestyle (reviewed in (68)). Indeed, *Pantoea* species are frequently associated with plant (or insect) hosts, however, the mode of *Pantoea* transmission along the host life-cycle remains to be clarified and reports of isolates from diverse soil habitats (13) indicate that *Pantoea* is well adapted to host-associated as well as free-living lifestyles. The LPP and pPag1 domestication most likely contributed to the versatility of *Pantoea* niche occupation.

Both LPP and pPag1 are related to plasmid lineages that are characterized by medium to large size and a gene content enriched in metabolic functions, similarly to other plasmids that are prevalent in plant microbiomes (69). Both LPP and pPag1 do not harbor homologs of *mob* relaxes, hence they are unlikely to be mobile. Nonetheless, the absence of plasmid mobility mechanisms does not preclude an evolutionary origin of these plasmids from a mobile ancestor, as plasmid mobility may be lost in the evolution of domesticated plasmids (70, 71). Plasmid domestication following the loss of plasmid mobility is bound to generate genetic polymorphism within the population, comprising plasmid-hosts and non-hosts sub-populations, that is the initial step of sympatric speciation. Evolution of stable plasmid inheritance will preclude the emergence of non-hosts from the host population, while the loss of plasmid mobility prevents gene flow from hosts to the non-hosts sub-population. The emergence of species isolation (termed also ‘reproductive barriers’) may be further promoted by selective sweep for the plasmid presence (i.e., for a trait supplied by the plasmid) and the consequent elimination of non-hosts from the population. Alternatively, migration of the hosts sub-population to a new niche made accessible by the plasmid presence may likewise lead to the emergence of new species. In the absence of genetic mechanisms for the translocation of beneficial plasmid genes to the host chromosome (e.g., transposable elements), domesticated plasmids are expected to become an integral component of the host genome in the form of chromids.

## Methods

### Genomes dataset

Genome assemblies of draft and complete genomes of *Pantoea* isolates were initially downloaded from RefSeq (72) (version of 03/2021). Additional *P. dispersa* genomes were downloaded from NCBI *a posteriori*. The genomes of insect symbionts were excluded as their genomes underwent extreme genome reduction (73). Such fast-evolving genomes may hinder the reconstruction of the species phylogeny, therefore we decided to restrict our study to plant-associated species (74). Additionally, we only analyzed classified species for which there is at least one finished genome. The final dataset comprises 33 complete and 220 draft *Pantoea* genomes. The complete list of genomes can be found in Table S1.

### Gene family reconstruction

For the reconstruction of gene families, the protein sequences of each possible pair of isolates was compared using blastp with an e-value ≤10^−6^ threshold (75). Reciprocal best blast hits are extracted and Needleman-Wunsch global alignments are computed with parasail using gap open and extension penalties of 11 and 1, respectively and the blosum62 matrix for amino-acid replacement (76). All pairs of sequences with ≥30% identical amino acids were clustered into gene families with MCL using an inflation index of 2 (77). The gene family reconstruction pipeline was used two times to reconstruct gene families among all *Pantoea* genomes, and among the *P. dispersa* genomes added *a posteriori*. The gene families of the two datasets were merged as follows: sequence similarity search among genomes of the two groups was performed and hits with a sequence similarity of ≥30% identical amino acids were retained. The average sequence similarity between each possible gene pair in a gene family was calculated. Gene family pairs with an average sequence similarity of ≥30% identical amino acids were retained and the best match were identified as the same gene family.

### Replicon assignment and plasmid classification

To identify and classify plasmid sequences, we compared the gene content across all contigs in the dataset. The overlap coefficient (78) in protein families encoded by each pair of contigs estimated as:

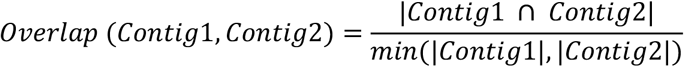

Contig pairs having ≥0.8 overlap coefficient and a length ratio ≤1/2 were retained for clustering. These thresholds were selected based on the minimum overlap found among 11 LPP plasmid and 36 chromosomal replicons (Fig. S11). Contig pairs were clustered using MCL with an inflation index of 2 (77). A total of 39 clusters including at least one plasmid identified as such in a completely sequenced genomes included plasmid-like 462 contigs. Most of these contigs (366 contigs, 80%) harbored one out of the two most common replication initiation (Rep) gene families in the dataset, indicating that the genus hosts mainly two major plasmid lineages. The largest plasmid lineage comprises sequences that were identified as the Large Pantoea Plasmid in previous studies (LPP; (29), followed by the pPag1 plasmid lineage. Rep gene family phylogenies were reconstructed to further validate the presence of such plasmid lineages (Fig. S2). In total, we identified 214 LPP-related contigs in 6 clusters of homologous plasmids and 114 pPag1-related contigs in 2 clusters of homologous plasmids. Clusters of homologous LPP were termed based on previous classification of LPP groups (29). Note that the LPP group III is not described in our dataset. Preliminary analyses including a larger number of isolates showed that the two strains reported to harbor this plasmid group (*Pantoea sp*. At-9b and *Pantoea cypripedii* LMG265^T^) are rather far relatives from the here studied taxa, and therefore we decided to exclude them from the current study. The remaining plasmid groups are either inconsistently present in the lineages or have a small number of contigs and therefore were not considered further in the analysis (Fig. S12). A plasmid network was reconstructed with Cytoscape v3.10.1 (79) using a threshold of overlap coefficient ≥ 0.4.

Chromosomal contigs were further validated as such using PlasClass (80), which calculates the probability of a contig to be a plasmid. We used contigs of known plasmids and chromosomes in the dataset as a training set and classified contigs as chromosomal when the probability of being a plasmid was ≤0.045. This threshold was chosen based on the probabilities inferred for chromosomes from completely sequenced genomes in the dataset. As a final quality check, we examined the distribution of chromosome size per isolate after assigning chromosomal sequences and compared it to the expected chromosomal genome size in each species (Fig. S13).

### Inference of species, plasmid and gene phylogenies

The *Pantoea* species tree was inferred from 299 single copy universal gene families that are exclusively found on the chromosome. Alignments for each gene family were reconstructed with MAFFT with default parameters (81). Maximum likelihood trees were inferred with IQ-TREE version 2.0.3, using the partitioned analysis (82). We used a general time reversible substitution model (GTR), using empirical base frequencies (+F). Gene trees were also reconstructed for replicon exclusive and replicon-shared gene families, using the same substitution model parameters. The root of the species tree was inferred by applying a phylogenomic rooting approach (35) using our dataset and, additionally, using the dataset provided by Crosby et al. (34) (see also Supplementary Note 1). The root inference of individual gene families was done with MAD (36). Isolates branching within the nine clades corresponding to nine main *Pantoea* species were classified as the corresponding species.

Plasmid trees were inferred using a subset of isolates to include the maximum number of complete single copy (CSC) gene families as described above for the species tree.

### Verticality tests

Replicon-specific single-copy gene families that comprise at least four sequences from at least two species were tested for deviation from the species phylogeny. A constrained topology was generated according to the species tree, where the sequences are split into the nine studied species. Multiple genes within the same species were added as a polytomic group. A constrained species trees was adapted to each gene family by pruning the species tree to contain only the sequences found in a focal gene family using ETE3 Toolkit (83). For each gene family, two phylogenies are inferred: an unconstrained maximum-likelihood phylogeny and a constrained-topology phylogeny using IQ-TREE version 2.0.3 (82, 84). The likelihood of the resulting topologies was compared using the Kishino-Hasegawa (KH) test using IQ-TREE. If the likelihood of the two phylogenies were not significantly different, the gene family is concluded to have evolved vertically.

### *Pantoea agglomerans* strain R1, growth conditions and marker frequency analysis

The *P. agglomerans* strain R1 is a member of the native, seed-borne microbiota of *Triticum aestivum* that was isolated from surface-sterilized seeds of the winter wheat cultivar Runal harvested in 2017 at the experimental farm Hohenschulen of Kiel University (85). The strain was routinely grown in Reasoner’s 2A medium at 30 °C. Whole genome DNA preparations (gDNA) for sequencing were prepared from either stationary-phase overnight cultures or from cells harvested during early logarithmic growth phase at OD450 ∼ 0.16 using the Wizard® Genomic DNA Purification Kit (Promega). For MFA, for each of the three sequenced replicates of exponentially growing and stationary cells, gDNA was isolated from independently grown cultures (30 ml) inoculated with a single colony. Sequencing using paired-end libraries with an insert size of 350-bp was performed on the Illumina NovaSeq platform with 150-bp read length on each end by NovoGene (Munich, Germany). Samples were sequenced to an average depth of ca. x250. The reads were adapter- and quality-trimmed with BBDuk from the BBTools package (BBMap – Bushnell B. – sourceforge.net/projects/bbmap/) using k-mer size of 23 and quality-trimming with a minimum phred score of 10. The reads were aligned to the circular scaffold of the chromosome of the strain R1, for which the contigs (NZ_WSSS01000001 to NZ_WSSS01000008) were sorted with MAUVE (86) using the genomes the of *P. agglomerans* strains DBM3797 (GenBank: CP086133.1) and L15 (GenBank: CP034148.1) as reference. The contigs of LPP and pPag1 represent the complete plasmids (confirmed by Sanger sequencing of PCR products bridging the contig junctions). The marker frequency analysis was performed using the MFA-seq protocol (87), which uses the localMapper bash script for aligning reads and calculating the sequencing depth in genomic windows, and the R-package Repliscope version 1.1.1 (88) for normalization.

### Functional annotation

Functional annotation and COG classification of the shared genes between chromosomes and LPP plasmids found among the set of 13 representative genomes (Table S1) were performed with eggNOG-mapper version 2 (89). Briefly, we mapped each gene sequence to the eggNOG database. When annotating gene families, the most common category or categories in each gene family was identified as the COG category for the whole gene family. The majority rule was not applied when annotating individual genes (i.e., in the species pangenome analyses).

### Statistical comparison of rooted tree topologies

To test for statistical support of the “translocation” and “redundancy” hypotheses, we rooted gene trees in Fig. 4 using MAD (36). The percentage of gene families within each scenario (“early” and “late”) that support each candidate root was calculated (Table S4, Fig. S1). Gene families included in “early” scenarios feature PAPs 1 and 2, while genes families included in “late” scenarios feature PAPs 3 and 4 as shown in Fig. 4a. The tested candidate roots comprise the set of inferred roots found among the complete set of genes (following the phylogenomic rooting approach (35)). These roots were also inferred among gene families in the complete dataset and are depicted in Fig. S1. The inferred root in a gene family corresponds to the candidate root with the minimal ancestral deviation (AD). In gene families where the ratio between the best root and the second-best root (Ambiguity Index) is equal or larger than 0.9, the gene family was concluded to support each root with half their weight.

### Phylogenetic pairwise distance analysis

The distribution of phylogenetic distances was examined between sequence pairs belonging to 617 complete single copy (CSC) gene families shared between the plasmid (LPP or pPag1) and the chromosome in the set of 13 representative genomes (Table S1). Only families having three or more homologs were included. For matter of comparison, distances were calculated also for chromosomal complete single-copy gene families (CSCs). The phylogenetic distance was calculated as the sum of branch length between homologous gene pairs (i.e., OTUs) in family gene trees inferred using Maximum Likelihood and the GTR+F evolutionary model using IQ-TREE version 2.0.3 (82).

## Data availability

Sequence reads are available at the European Nucleotide Archive (ENA; accession: PRJEB81354). All other material is supplied in the supplementary information.

## Acknowledgements

We thank Giddy Landan for insightful comments on the phylogenetic analyses. We thank Kai Ripcke and Shreya Vichare for fruitful discussions and Yiqing Wang, Fabian Nies and Dustin Hanke for critical comments on the manuscript. This work was supported by the DFG funded CRC1182 Origin and Function of Metaorganisms and ERC grant pMolEvol (grant number 101043835).

## Author contributions

DRP and TD designed the study; DRP collected the data and performed all comparative genomics and phylogenetic analyses; NFH and PK performed the marker free analysis (MFA). DRP and TD interpreted the results with contribution from NFH and PK. DRP and TD wrote the manuscript with contributions from all authors.

