## Supplementary Material for "Evolution of plasmid domestication in plant-associated *Pantoea*: massive gain of genetic redundancy followed by differential gene loss"

**This PDF file includes:**

Supplementary Note 1: Species phylogeny root inference
Figures S1 to S12
Table S2 to S5
SI References

**Supplementary Note 1: Species phylogeny root inference**

To identify the best root candidate among chromosomal gene trees we used the phylogenomic rooting approach with the dataset as in Fig. 1 (1). Briefly, root candidate partitions were identified among 299 complete single-copy chromosomal genes (CSCs) by collecting the best MAD candidate root partition per gene tree (2). Then, these partitions are mapped onto partial and multicopy genes, and the distribution of their ancestral deviations (AD) is analyzed. In total, 6,612 gene trees were included. This yielded to high ranking 8 candidate root partitions (Fig. S3). Candidate root partition 1, that splits *P. stewartii*, *P. allii* and *P. ananatis* from the remaining species appears most frequently as the best partition among CSC trees (64%), followed by the candidate root partition 2, that splits *P. eucrina* and *P. dispersa* from the remaining species (27%). Then, we performed pairwise root support tests to study whether the most frequent candidate root partition 1 among CSC trees is also the best partition when considering the AD distribution among all gene trees (Table S2). We did not find another candidate root partition significantly better than partitions 1 and 2. Additionally, there was no significant difference between candidate root partitions 1 and 2, and therefore, we identified them as competing root partitions.

To further investigate the root position, we repeated the analysis using an extended dataset that includes 425 genomes and 50 species provided by Crosby et al. (3). Here, we identified and removed potential LPP plasmids from the dataset using fastANI (4). Briefly, we calculated ANI (average nucleotide identities) between the contigs in the Crosby et al. dataset and the contigs identified as LPP in this study. Contigs with ANI over 80% and 80% coverage were identified as LPP plasmids. This yielded 329 genomes, most of which include a single LPP contig (in the remaining genomes the LPP pans multiple contigs). Gene families identified among putative LPP contigs were discarded for the phylogenomic root analysis. In total, we analyzed 12,354 gene trees and identified 107 candidate root partitions among 205 CSC genes. We examined the AD distribution among the 13 best candidate root partitions (median CSCs AD $\leq$ 0.336). We performed pairwise root support tests and found that the best partition is significantly better than any other partition in the set (FDR adjusted p-value = 0.00016). This root partition coincides with the outgroup used in Crosby et al. (3), which corresponds to the split between *P. eucrina* and *P. dispersa* and the remaining species in our dataset. Therefore, we conclude that the better supported root partition in our dataset splits *P. dispersa* and *P. eucrina* from the remaining species.

### Supplementary Figures

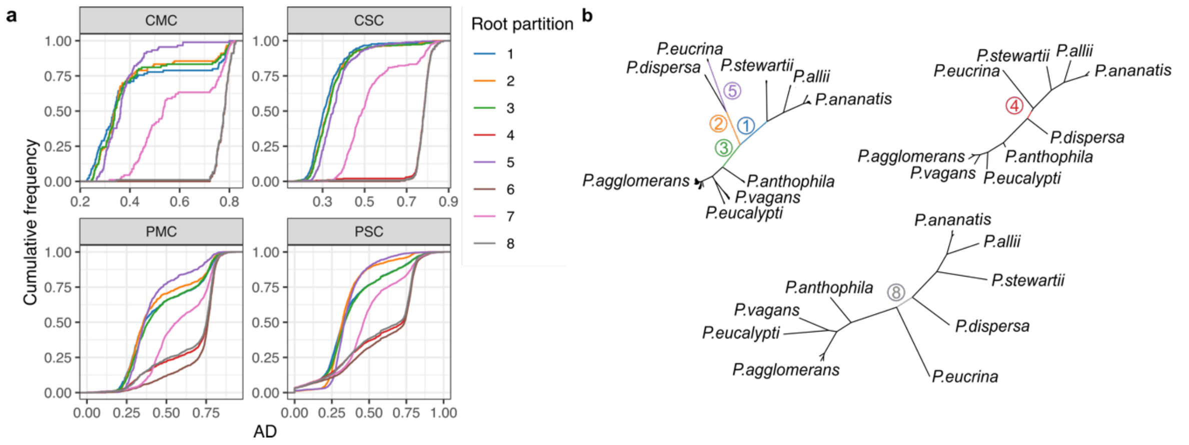


**Supplementary Figure S1. Phylogenomic rooting of chromosomal genes for 253 *Pantoea* genomes. a,** Ancestral deviation (AD) distributions for 299 complete single-copy gene trees (CSC), 90 complete multi-copy gene trees (CMC), 5,011 partial single-copy gene trees (PSC) and 1,212 partial multi-copy gene trees (PMC). **b,** Root partitions found in (**a**). Root partitions 6 and 7 are not represented in the trees as they were only found as best candidate root partitions in one gene tree each. Additionally, the two trees where these partitions are found are highly polytomic and species do not form defined clusters. Partition 6 splits *P. dispersa*, *P. ananatis*, *P. allii* and *P. eucrina* from the remaining species. Partition 7 splits *P. eucalypti*, *P. vagans* and *P. agglomerans* from the remaining species.

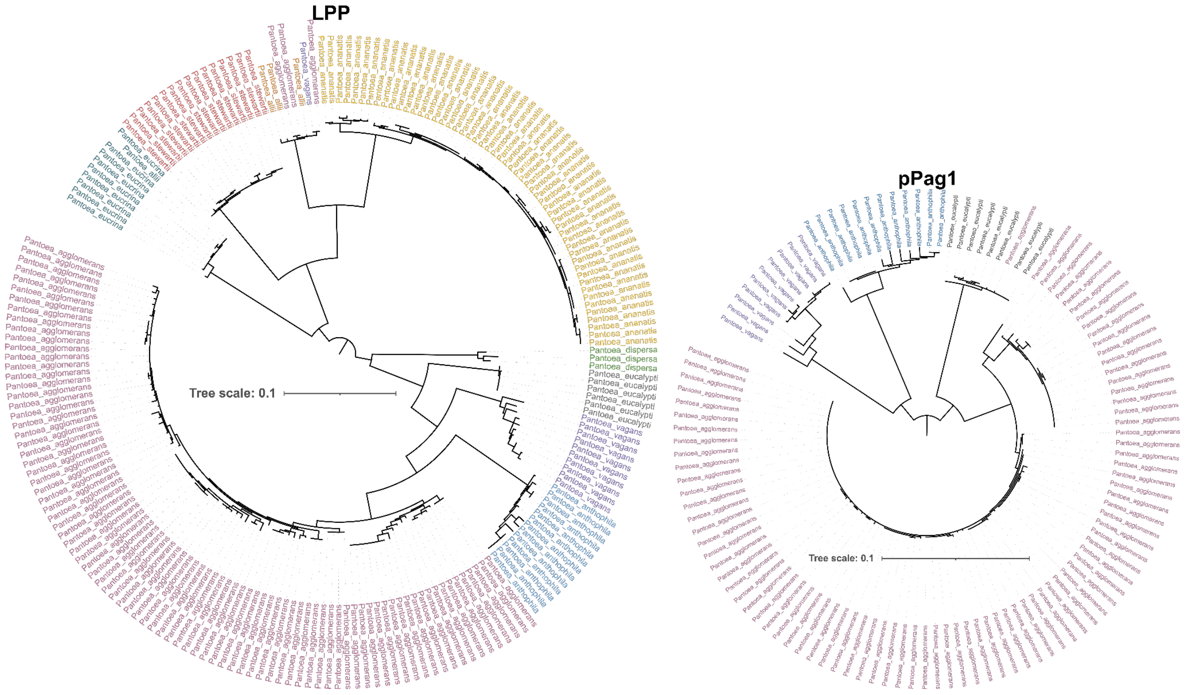


**Supplementary Figure S2. LPP and pPag1 replication initiation (*repB*) gene family phylogenies.** Maximum Likelihood phylogenies inferred using a general (GTR+F) model of evolution using IQ-TREE (5). The root was inferred by midpoint rooting. Species labels correspond to the main identified species in this study.

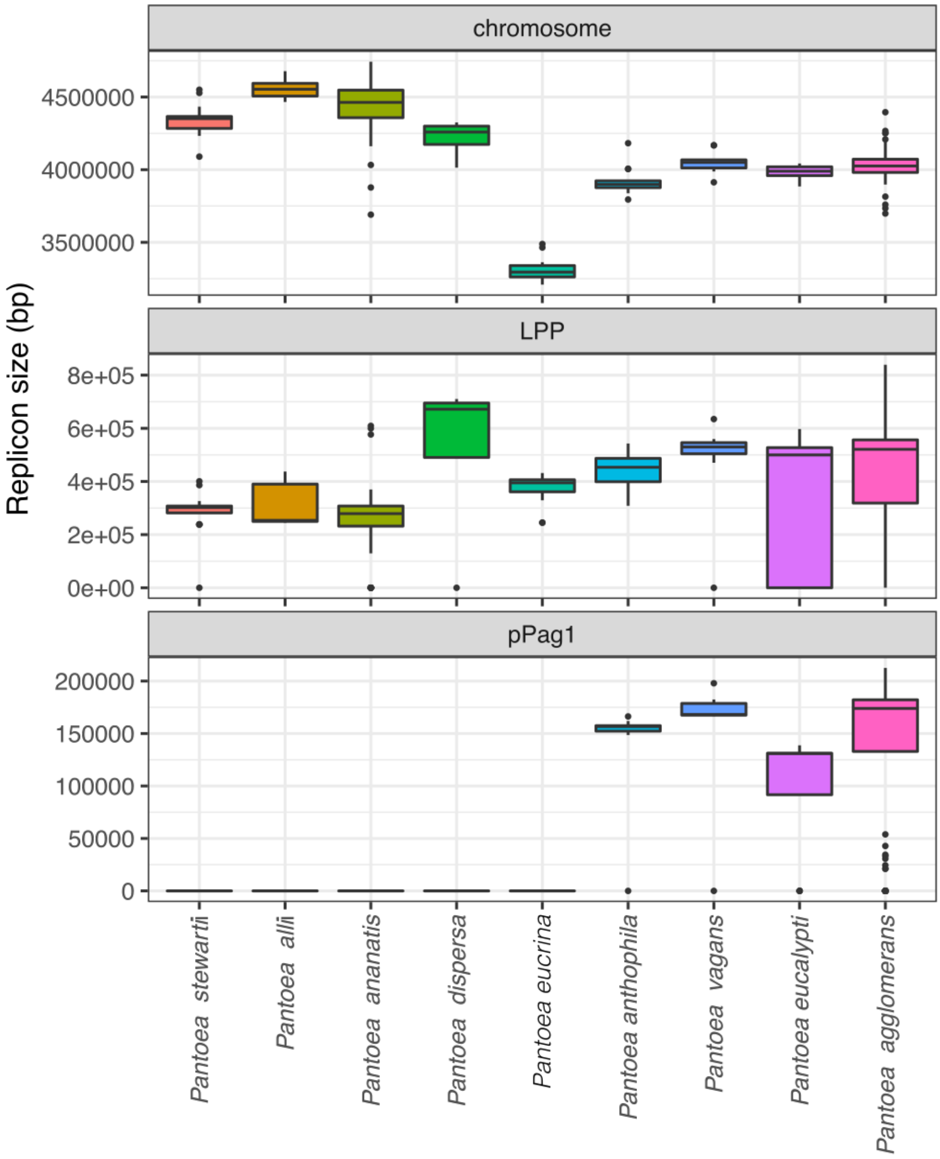


**Supplementary Figure S3. Distribution of replicon size per species.**

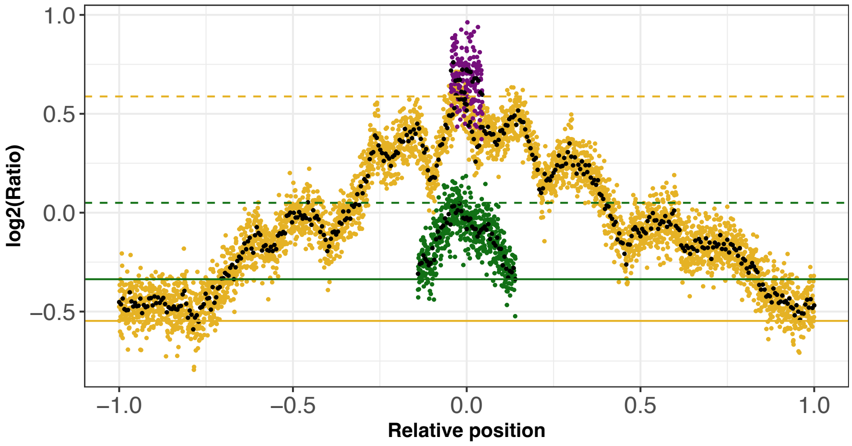


**Supplementary Figure S4. Marker frequency analysis (MFA) for LPP (green), pPag1 (purple) and chromosome (yellow).** This figure presents the results of a second, independent, replicate of the MFA analysis. The number of reads is presented for 1 kb (colored dots) and 10kb (black dots) windows as the $log_2$ ratio of the number of reads divided by the number of reads of a non-replicating sample. Additionally, the number of reads is normalized by the total number of reads per sample. The x-axis indicates the relative position on the chromosome, where 0 corresponds to the origin of replication in all three replicons, and 1 and -1 correspond to the replication termination in the chromosome. Dashed lines indicate the origin of replication while continuous lines indicate termination in LPP and pPag1.

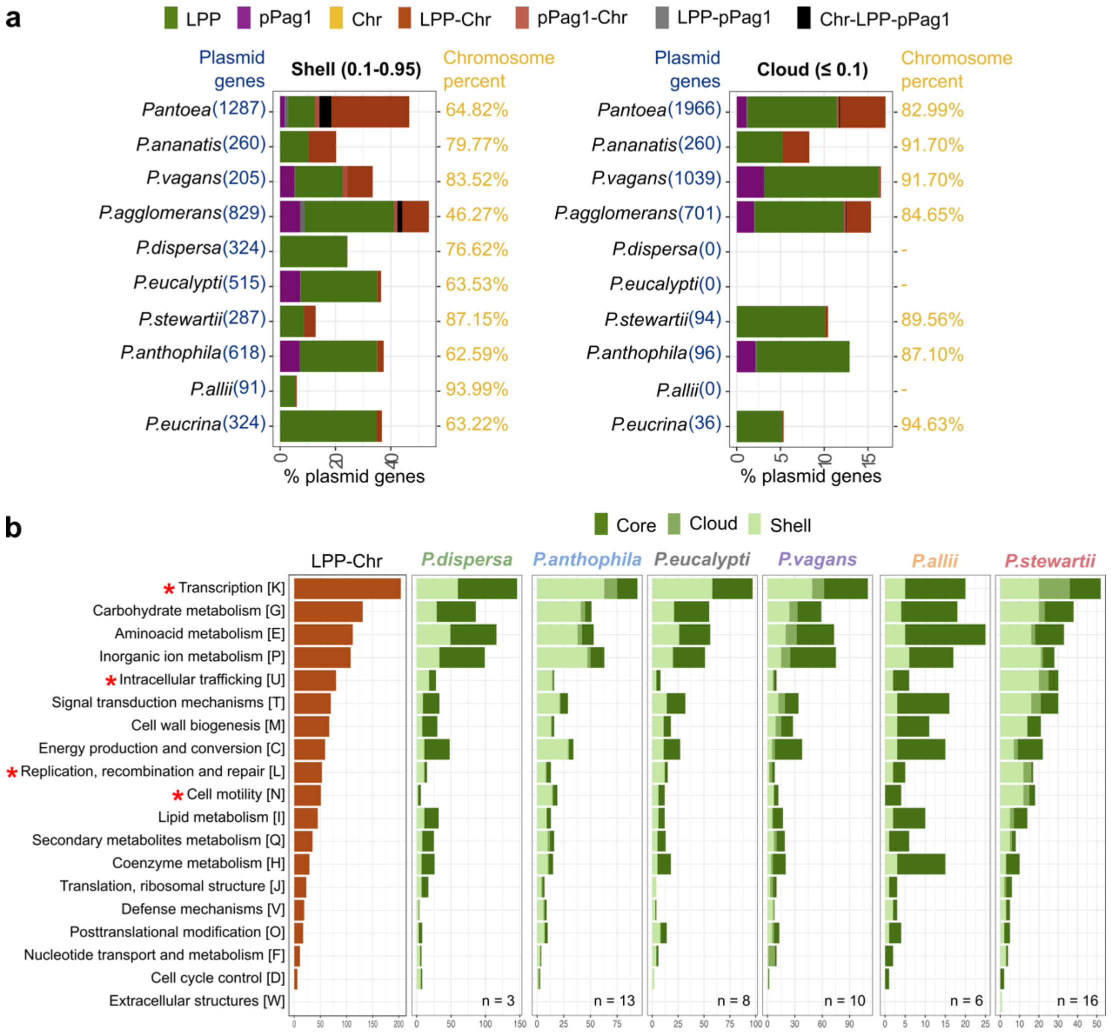


**Supplementary Figure S5. Gene families shared between the chromosome and pPag1 in the LPP pangenome. a,** Distribution of shell (frequency between 0.1 and 0.95) and cloud genes (frequency equal or smaller than 0.1) in the *Pantoea* pangenome in 211 isolates harboring the LPP plasmid. Replicon location of genes is shown in different colors. The genome of *P. dispersa* stands out as all shell gene families are exclusively found in either the chromosome or the LPP. Note that, the pangenome shape depends on the number of genomes sampled per species. **b,** Functional composition of shared genes between LPP and chromosome and the LPP pangenome per species not shown in main Fig. 3. Sample size (n) shows the number of isolates per species.

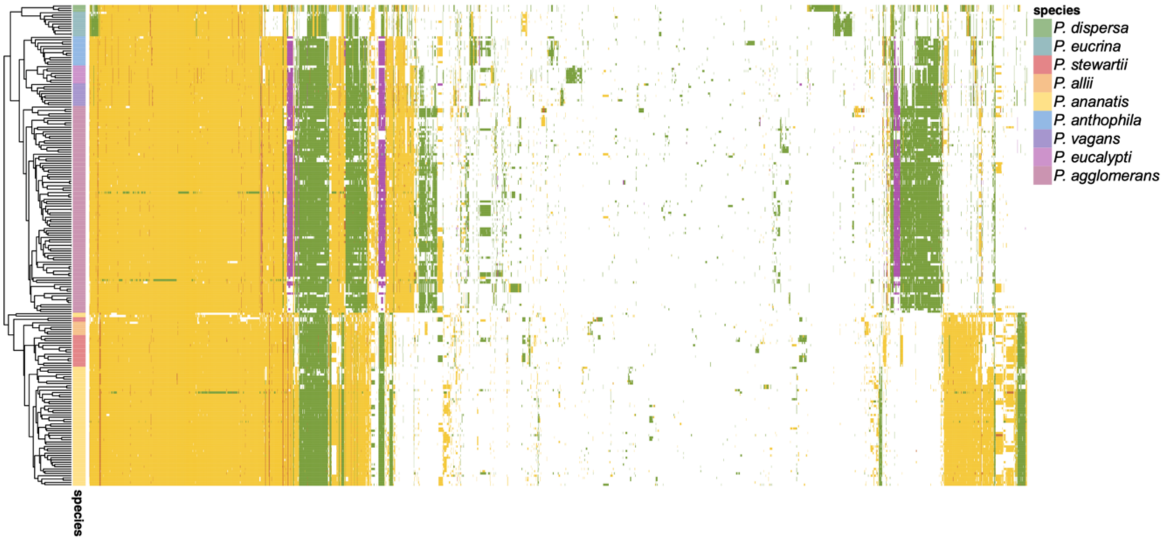


**Supplementary Figure S6. Presence, absence and genomic location of gene families from the LPP pangenome in 214 isolates**. Note that two *P. agglomerans* and one *P. ananatis* isolates present many LPP encoded genes which appear to be found on the chromosome for the remaining isolates of the species. This aberrant pattern suggests possible assembly artifacts.

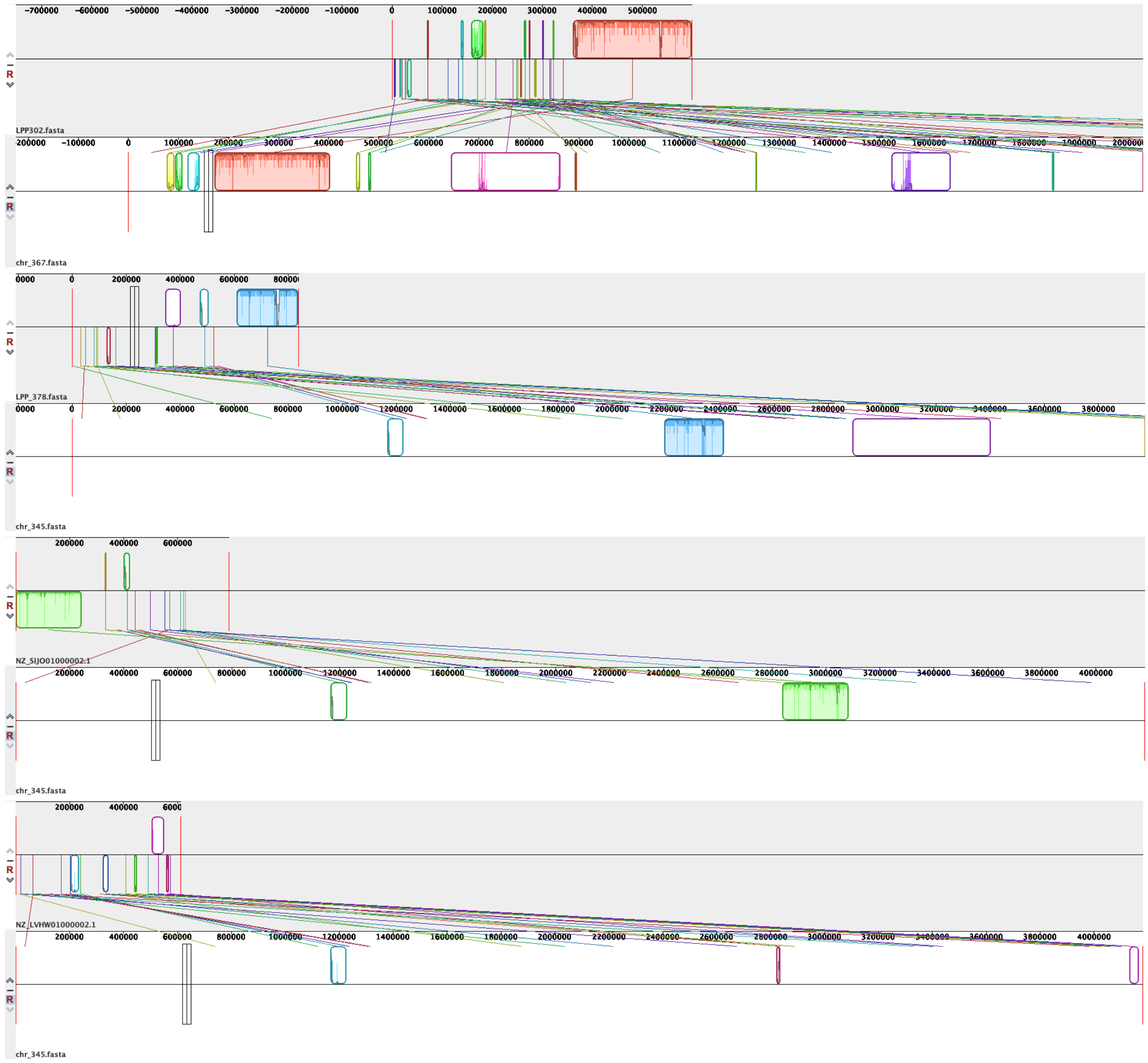


**Supplementary Figure S7. Pairwise alignments between LPP from excluded isolates and the chromosome of representative isolates from the same species.** The first alignment belongs to *P. ananatis* and the next two alignments to *P. agglomerans* isolates. The figure below corresponds to the alignment for a non-excluded pair of isolates for comparison. Solid blocks correspond to high similarity regions between the plasmid and the chromosome. These blocks are typically allocated on the edges of the plasmid contigs. Additionally, such high similarity blocks are absent in non-excluded isolates. This supports our conclusion that the LPP contigs from the three first isolates likely are the result of a mis-assembly. The alignments were performed with Mauve (6).

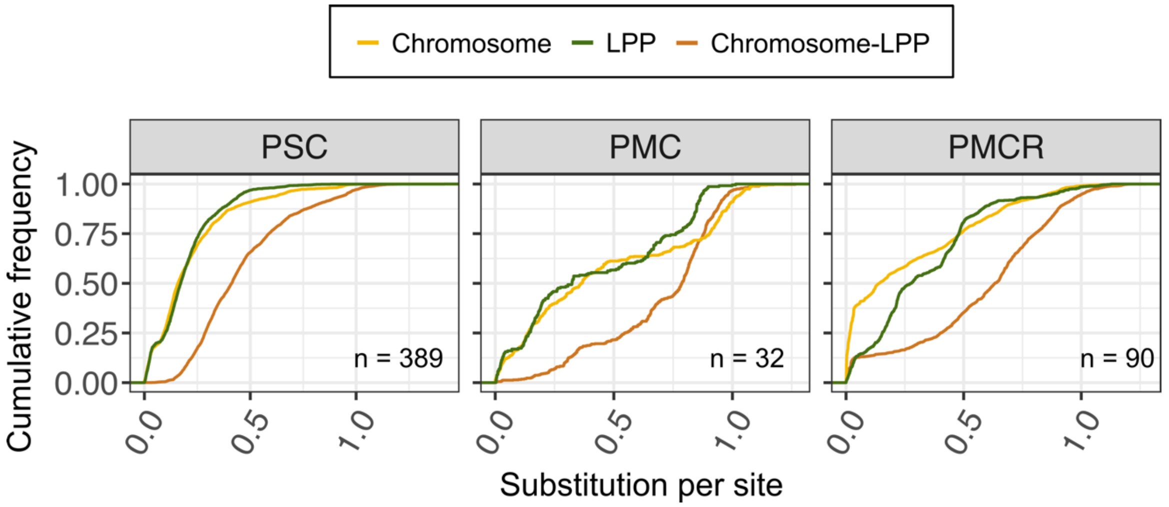


**Supplementary Figure S8. Pairwise distances in partial single- and multi- copy gene families.** Distribution of pairwise phylogenetic distance calculated as substitution per site across partial gene families shared between chromosome and LPP plasmid. The dataset comprises partial single-copy gene families (PSC); partial multicopy gene families, where copies are present in a single replicon (PMC); partial multicopy gene families, where copies can be present in both replicons (MCR). (n) shows the number of gene families analyzed.

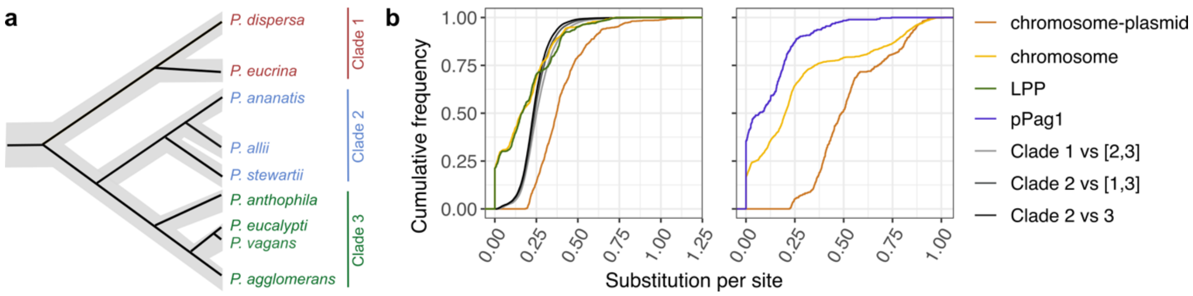


**Supplementary Figure S9. Distribution of pairwise phylogenetic distance calculated as substitution per site across gene families shared between the chromosome and the LPP. a,** Main phyletic clades that differ in the location of replicon-shared genes. **b,** Phylogenetic distance distribution between the main phyletic groups that split chromosome and plasmid sequences in the main evolutionary scenarios, compared to CSCs shared between chromosome and LPP (see main Fig. 4d). The dataset comprises 2,071 of CSC genes found exclusively in the chromosome.

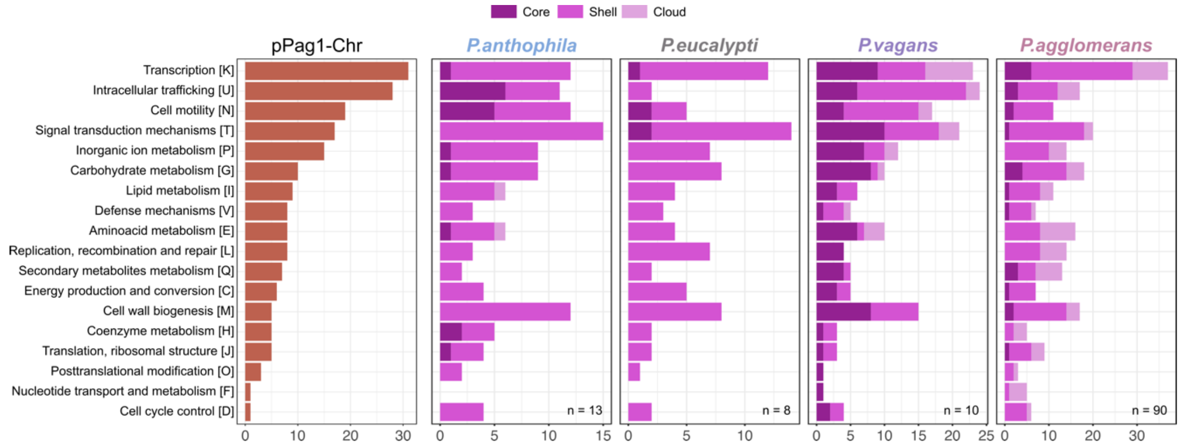


**Supplementary Figure S10. Functional composition of pPag1 gene families.** Functional category distribution of shared genes between pPag1 and chromosome and genes of the pPag1 pangenome per species.

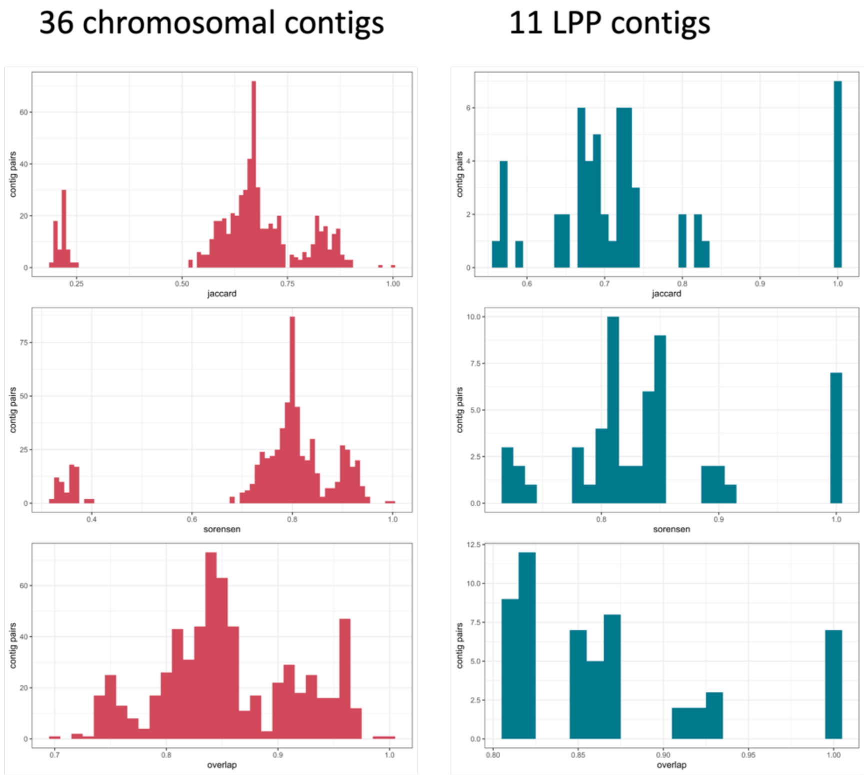


**Supplementary Figure S11. Measures of gene content similarity.** The distribution of gene content similarity using different measures among known chromosome and LPP plasmid sequences in complete genomes from the dataset.

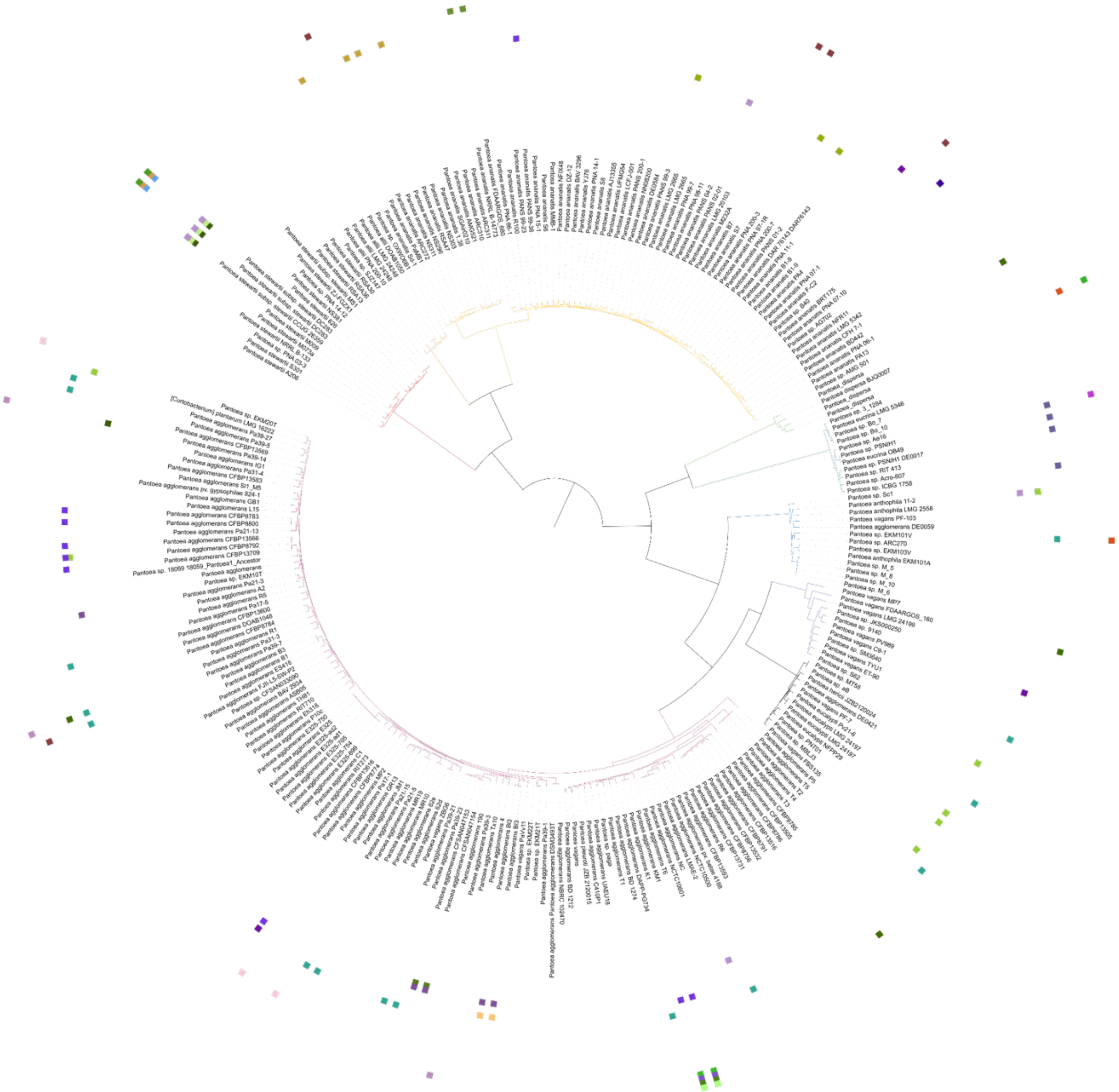

**Supplementary Figure S12. Distribution of non-LPP and non-pPag1 plasmids in the dataset.** Each outer ring represents a different plasmid group.

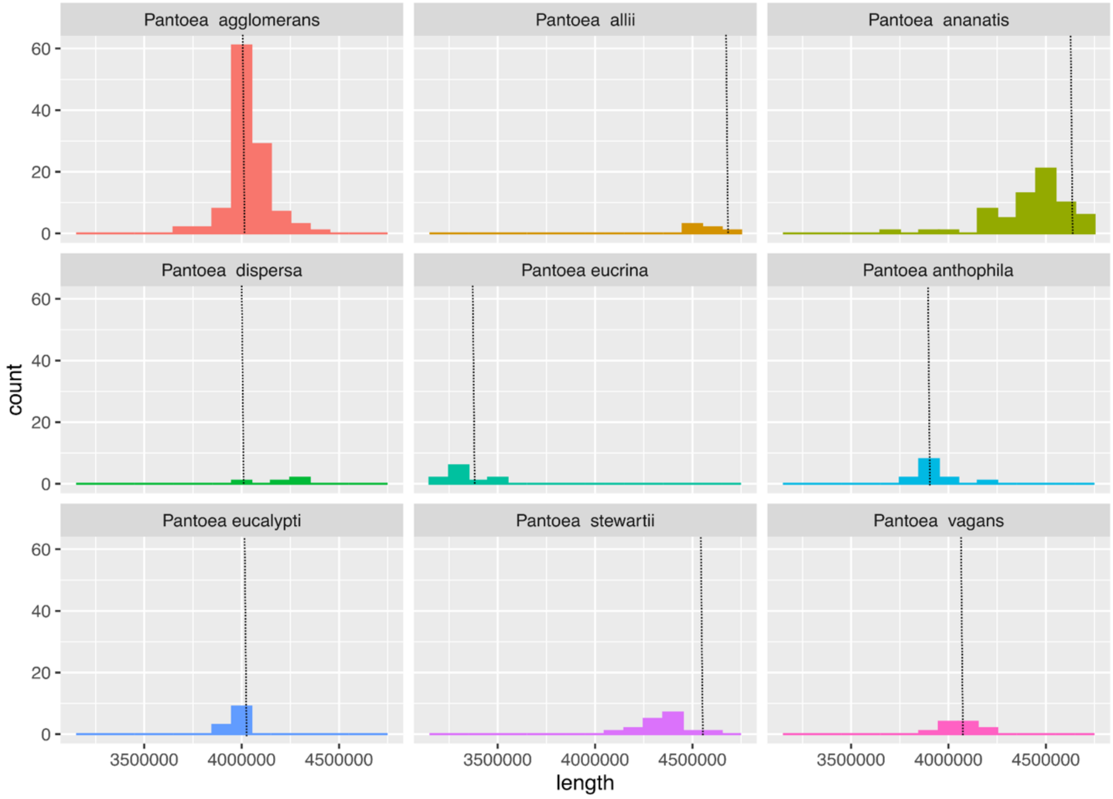


**Supplementary Figure S13. Chromosome size distribution per species.** Dashed lines indicate the expected chromosome size for representative genomes in each species. (Suppl. Table 1)

### Supplementary Tables

**Supplementary Table S1.** Genomes dataset is available as an Excel sheet (Table_S1.xlsx).

**Supplementary Table S2. Pairwise root partition support tests.** Values are FDR adjusted p-values of the two-sided Wilcoxon signed-rank test. Red highlights indicate competing root positions ($\alpha$ = 0.05). Rows and column names correspond to the partitions shown in Fig. S1.

| | 1 | 2 | 3 | 4 | 5 | 6 | 7 |
|---|---|---|---|---|---|---|---|
| 2 | 0.51799 | | | | | | |
| 3 | 0.00018 | $7.5x10^{-7}$ | | | | | |
| 4 | $< 2x10^{-16}$ | $< 2x10^{-16}$ | $< 2x10^{-16}$ | | | | |
| 5 | 0.00029 | $4.1x10^{-8}$ | 0.47454 | $< 2x10^{-16}$ | | | |
| 6 | $< 2x10^{-16}$ | $< 2x10^{-16}$ | $< 2x10^{-16}$ | $1.1x10^{-6}$ | $< 2x10^{-16}$ | | |
| 7 | $< 2x10^{-16}$ | $< 2x10^{-16}$ | $< 2x10^{-16}$ | $< 2x10^{-16}$ | $< 2x10^{-16}$ | $< 2x10^{-16}$ | |
| 8 | $< 2x10^{-16}$ | $< 2x10^{-16}$ | $< 2x10^{-16}$ | 0.22145 | $< 2x10^{-16}$ | $2.9x10^{-10}$ | $< 2x10^{-16}$ |

**Supplementary Table S3. Distribution of COG functional categories across species for LPP and pPag1 pangenomes.** Values are FDR adjusted p-values of the Pearson's Chi-squared tests. Red highlights indicate functional categories that are under or overrepresented across species (FDR < 0.05).

| COG category | p-value (LPP) | p-value (pPag1) |
|---|---|---|
| C | $9.28x10^{-1}$ | $9.86x10^{-1}$ |
| D | $9.50x10^{-1}$ | $9.86x10^{-1}$ |
| E | $2.76x10^{-1}$ | $9.86x10^{-1}$ |
| F | $9.28x10^{-1}$ | $5.89x10^{-1}$ |
| G | $9.28x10^{-1}$ | $9.86x10^{-1}$ |
| H | $2.84x10^{-1}$ | $9.86x10^{-1}$ |
| I | $7.25x10^{-1}$ | $9.86x10^{-1}$ |
| J | $9.28x10^{-1}$ | $9.86x10^{-1}$ |
| K | $1.37x10^{-2}$ | $9.86x10^{-1}$ |
| L | $2.33x10^{-2}$ | $5.89x10^{-1}$ |
| M | $5.02x10^{-1}$ | $9.86x10^{-1}$ |
| N | $2.30x10^{-4}$ | $5.89x10^{-1}$ |
| O | $4.47x10^{-1}$ | $9.86x10^{-1}$ |
| P | $5.28x10^{-2}$ | $9.86x10^{-1}$ |
| Q | $9.71x10^{-1}$ | $7.14x10^{-1}$ |
| T | $2.76x10^{-1}$ | $8.77x10^{-1}$ |
| U | $5.58x10^{-5}$ | $1.86x10^{-1}$ |
| V | $5.14x10^{-1}$ | $9.86x10^{-1}$ |
| W | $3.59x10^{-1}$ | - |

**Supplementary Table S4. Percentage of genes shared between chromosome and LPP plasmid supporting each candidate root position.** The results are shown separately for each evolutionary scenario. Candidate root ids correspond to those shown in Fig. S1. Genes included in "Early scenarios" comprise those in PAPs 1, 2 from Fig. 4a. Genes included in "Late scenarios" comprise those in PAPs 3, 4 from Fig. 4a. Additionally, we performed the analysis in PAP 5 from Fig. 4a. Highlighted in red is the most supported candidate root position in each scenario. Note that one gene may support several scenarios, therefore contributing partially to each scenario.

| Candidate root | Early scenarios | Late scenarios | PAP 5 |
|---|---|---|---|
| 1 | 22% | 52% | 6% |
| 2 | 48% | 21% | 31% |
| 3 | 18% | 4% | 40% |
| 4 | 4% | 8% | 10% |
| 8 | 8% | 15% | 12% |
| Number of genes | 115.5 | 76 | 24.5 |

**Supplementary Table S5. Average phylogenetic distance within and between replicons for gene families shared between chromosome and LPP.** The dataset comprises complete single-copy gene families (CSC); complete multicopy gene families, where copies are present in a single replicon (CMC); complete multicopy gene families, where copies can be present in both replicons (CMCR); partial single copy gene families (PSC); partial multicopy gene families, where copies are present in a single replicon (PMC) and partial multicopy gene families, where copies can be present in both replicons (PMCR).

| Gene group | Replicon | Median(±IQR) |
|---|---|---|
| CSC | Chr-LPP | 0.372(±0.178) |
| | Chromosome | 0.229(±0.203) |
| | LPP | 0.218(±0.228) |
| CMC | Chr-LPP | 0.47(±0.205) |
| | Chromosome | 0.215(±0.149) |
| | LPP | 0.351(±0.233) |
| CMCR | Chr-LPP | 0.47(±0.205) |
| | Chromosome | 0.215(±0.149) |
| | LPP | 0.351(±0.233) |
| PSC | Chr-LPP | 0.412(±0.302) |
| | Chromosome | 0.158(±0.201) |
| | LPP | 0.172(±0.161) |
| PMC | Chr-LPP | 0.787(±0.332) |
| | Chromosome | 0.362(±0.763) |
| | LPP | 0.324(±0.618) |
| PMCR | Chr-LPP | 0.632(±0.402) |
| | Chromosome | 0.141(±0.469) |
| | LPP | 0.276(±0.325) |
